# BOLD response is more than just magnitude: improving detection sensitivity through capturing hemodynamic profiles

**DOI:** 10.1101/2023.02.13.528362

**Authors:** Gang Chen, Paul A. Taylor, Richard C. Reynolds, Ellen Leibenluft, Daniel S. Pine, Melissa A. Brotman, David Pagliaccio, Simone P. Haller

## Abstract

Typical FMRI analyses assume a canonical hemodynamic response function (HRF) with a focus on the overshoot peak height, while other morphological aspects are largely ignored. Thus, in most reported analyses, the overall effect is reduced from a curve to a single scalar. Here, we adopt a data-driven approach to HRF estimation at the whole-brain voxel level, without assuming a response profile at the individual level. Then, we estimate the response in its entirety with a roughness penalty at the population level to improve predictive accuracy, inferential efficiency, and cross-study reproducibility. Using a fast event-related FMRI dataset, we demonstrate the extent of under-fitting and information loss that occurs when adopting the canonical approach. We also address the following questions:

1. How much does the HRF shape vary across regions, conditions, and groups?
2. Does an agnostic approach improve sensitivity to detect an effect compared to an assumed HRF?
3. Can examining HRF shape help validate the presence of an effect complementing statistical evidence?
4. Could the HRF shape provide evidence for whole-brain BOLD response during a simple task?

## 1 Introduction

The blood-oxygen-level-dependent (BOLD) signal, acquired via functional magnetic resonance imaging (FMRI), is considered a proxy for neuronal activity. Neurons are constantly in need of oxygen and glucose to maintain their functioning. In particularly demanding contexts (e.g. performing a cognitively intensive task), energy-rich, oxygenated blood oversupplies active regions. This neurovascular coupling also plays a role in supplying oxygen for neuromodulator synthesis, signaling to neurons, and other physiological functions (Drew, 2022). Thus, the hemodynamic response function (HRF) profiled by the BOLD signal in FMRI serves as an indirect indicator for neuronal activity.

The prototypical HRF shape has been broadly verified, yet the mechanisms that generate it remain elusive (Buxton, 2021). Briefly, BOLD signal relies on the measurement of T2*^*^* relaxation, which is sensitive to local concentrations of paramagnetic deoxyhemoglobin. It is generally recognized that, as a nonlinear function of neuronal activity, the BOLD response to an instantaneous stimulus consists of three phases relative to a baseline (Fig. 1A). First, a small and short downward change (referred to as the “initial dip”), which may immediately follow stimulus onset and last for as short as one second (Hong and Zafar, 2018). This dip is likely related to localized parenchymal increases of total hemoglobin prior to the dilation of pial arteries (Hillman, 2014) and is sometimes thought of as a direct marker of neuronal oxygen consumption. Second, a large “overshoot” that corresponds to neuronal activity with a very brief duration (Birn et al., 2001) and is usually the modeling focus (Logothetis et al., 2001). This primary positive BOLD response is directly associated with the properties of the brain’s vasculature (e.g., vasodilation) and neurovascular coupling that drive stimulus-evoked functional hyperemia. Finally, an “undershoot”^1^, which may last up to 30 seconds (Yacoub et al., 2006; Poser et al., 2010). This negative BOLD response may be partly caused by neural suppression, a residual high concentration of total hemoglobin in the capillary beds and decreased metabolic rate of oxygen consumption, combined with frequently observed post-stimulus arterial vasoconstriction (Hillman, 2014).

**Figure 1:**
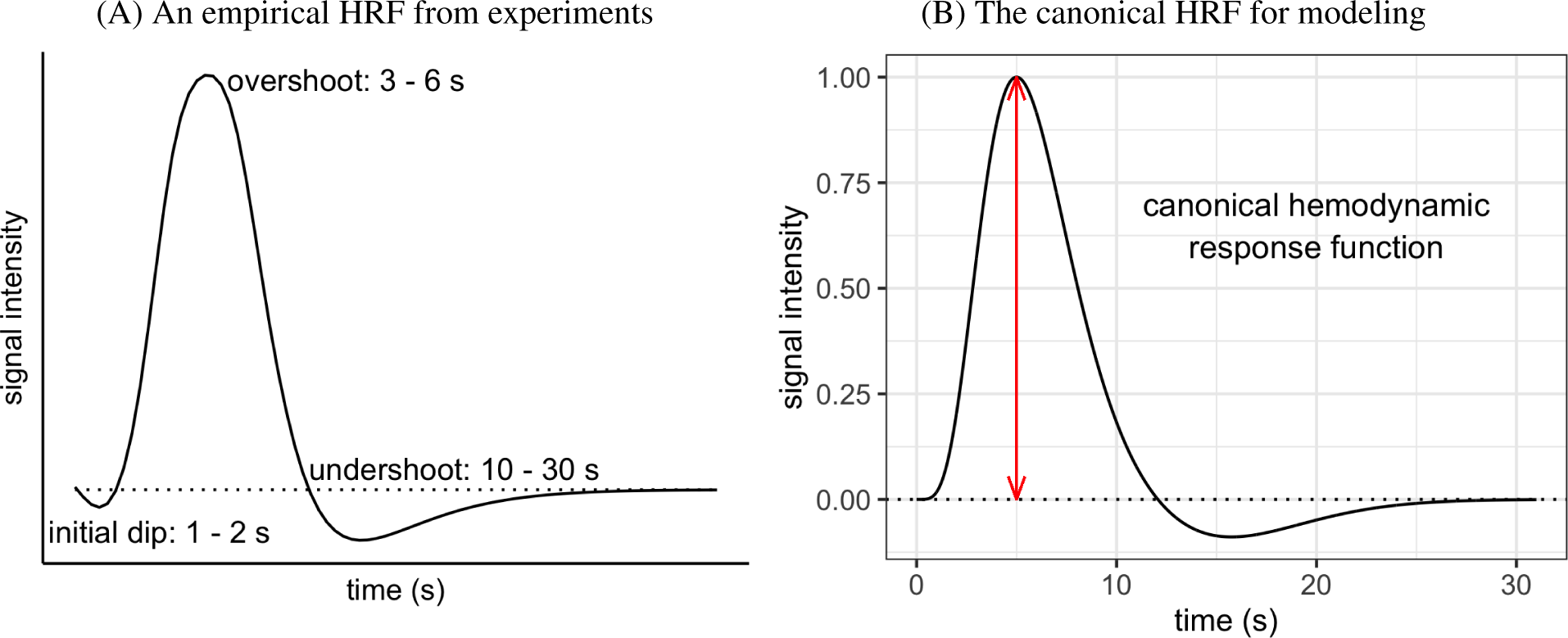
(A) Typical depiction of HRF shape and features based on FMRI and optical imaging experiments. The scale of the response intensity (vertical axis) depends on stimulus type, brain region, and local vasculature. (B) A canonical HRF for modeling (Friston et al., 1998) is usually constructed through an impulse response curve as a function of time *t*: here, *h*(*t*) = 5.7*t*^5^*e^−t^/*Γ(6) *−* 0.95*t*^15^*e^−t^/*Γ(16). It serves as a basis function and is convolved with stimulus timings to generate a series of expected HRFs for a task condition. These convolved HRFs are then used as regressors in a time series regression for acquired BOLD signals at the individual level. The overshoot peak occurs at 5 s and is scaled to 1 (red line) here. It is the sole focus in standard modeling, and the corresponding regression coefficient for a task condition with instantaneous trials can be conveniently interpreted as, for example, percent signal change. Moreover, the onset and duration of the undershoot are fixed, with a preset nadir of about 9% of the overshoot peak. The dotted horizontal line in both panels shows the baseline level of BOLD signal.

### 1.1 Canonical modeling approaches to capturing BOLD response and their pitfalls

The BOLD signal is measured over time in intervals of repetition time (TR). At the individual level, a regressor for each task condition is generated by convolving the associated stimulus timing with a canonical HRF under the assumption that the basic shape is fixed. The regressor is then entered into a time-series regression model and plays the role of “pattern matching” in the data. A canonical HRF is informed by biophysical experiments, and is implemented as different versions of gamma variates (Cohen, 1997; Friston et al., 1998). For example, the canonical HRF in Fig. 1B has a total duration of 32 s with a fixed duration (12 s) and peak location (5 s) of the overshoot, the nadir (*≈* 9% of the overshoot height) and duration (20 s) of the undershoot. In addition, no initial dip is contained in this basis shape. If a regressor associated with the canonical HRF is scaled to have a peak of one, then the corresponding regression coefficient can be conveniently interpreted as approximated BOLD response magnitude at the peak (e.g., in percent signal change, Chen et al. (2017)). The peak of the canonical HRF is always estimated to be at five seconds.

Accurate modeling with a fixed HRF depends on the relevant features being present at the right time (e.g., time to peak, relative magnitude of undershoot to peak, etc.) and meeting the assumption that the chosen shape features are constant across the the brain. It is well known that the HRF shape varies substantially across individuals, brain regions, tasks, individuals, and groups (by sex, age, diagnostic status) (Handwerker et al., 2004; West et al., 2019). This includes variability in peak timing, overshoot width, undershoot depth and duration, even with the same peak height. For instance, in an experiment with long blocks (e.g., 10 s or more), convolution with a canonical HRF would render a plateau in the model regressor - such a horizontal BOLD response is unlikely to be sustained due to habituation to the stimulus and other factors. Finally, the presence, size, and timing of the initial dip and undershoot can also vary substantially. Its characterization and the associated underlying complex mechanism remain a captivating research topic (e.g., Himmelberg et al., 2022; Drew, 2022; Taylor et al., 2022a). These aspects are currently not generally considered in modeling, even though they have been observed and they reflect physiological (and likely neuronal) response information. In addition, the undershoot is assumed to have a fixed depth relative to the preceding overshoot peak, notwithstanding that empirical data indicates there is no close relation (Aguirre et al., 1998). Although using a canonical HRF effectively reduces complexity in the modeling procedure, it seems unlikely that it would accurately represent the diversity and variability of actual responses; therefore, in many situations, it will under-fit the data, misrepresent the true physiology and miss useful information.

### 1.2 Ways to improve HRF modeling

Some potential improvements in modeling the response would be to adjust the canonical HRF or to extract multiple HRF features. For example, adding adaptive basis functions (Friston et al., 1998) can improve the overall fit by allowing variability of HRF profile features such as time-to-peak, full-width at half-maximum (FWHM), and peak duration. However, when these additional basis functions of temporal and dispersion derivatives are used, their associated effects are usually not examined at the population level; and even when included in individual-level modeling to improve fitting, most analyses still typically focus *only* on the peak magnitude associated with the canonical HRF at the population level, which ignores important information. Therefore, the potential for misrepresentation of effect estimates remains. Even when the effects associated with those specific profile features are incorporated into the population-level model, the potential gain in detection sensitivity is quite limited due to the subsequent information loss from information reduction before quantitative comparison (Lindquist et al., 2009; Chen et al., 2015). Moreover, HRF variation may occur beyond the profile features that have been considered. For example, the depth of the undershoot is generally fixed relative to that of overshoot in the canonical HRF (as shown in Fig. 1B), an a priori assumption that is likely often violated.

An potentially better approach is to directly estimate the HRF at each spatial unit (such as a voxel or ROI) from the data. Without assuming a prefixed shape, one could estimate HRFs, not through modeling, but by directly averaging the data across repeated instantiations using carefully designed experiments with well-separated trials or blocks (e.g., with an inter-stimulus interval of more than 10 s). While such a strategy has yielded interesting results in multiple studies (e.g., Handwerker et al., 2004; Gonzalez-Castillo et al., 2012), it is usually limited to exploratory investigations which just examine a few brain regions at the individual level and is not applied more broadly to typical experimental designs in the field (e.g., fast event-related experiments).

More broadly, instead of extracting a few limited HRF profile features, an agnostic strategy in modeling would be more precise through deconvolution. At the individual level, this can be achieved per spatial unit under assumptions of a certain BOLD response duration and a linear time-invariant system, using piecewise linear splines (tent, hat, triangular bases or sticks, Fig. 2A) or piecewise cubic splines in AFNI (Cox, 1996, or finite impulse response bases in FSL (Jenkinson et al., 2012) and SPM (Penny et al., 2011). Ultimately, the HRF is estimated at TR grids, each of which is associated with a basis or knot. Although these modeling approaches are maximally flexible in capturing the HRF shape, this flexibility comes at a cost: model complexity is substantially increased, as these approaches require many regressors per condition due to the number of bases used to cover the response duration. Moreover, the estimated HRFs are vulnerable to sampling fluctuations and could perform poorly when applied to out-of-sample data, a phenomenon with a long history motivating techniques of regularization, partial pooling, and hierarchical modeling (e.g., Stein, 1956; Efron and Morris, 1976; Hastie and Tibshirani, 1990; Wood, 2017). The estimated HRFs through deconvolution are only sampled at discrete time points. Therefore, simple linear interpolation would lead to a jagged appearance (Fig. 3A). Moreover, overfitting may occur and lead to compromised out-of-sample predictive accuracy when no restrictions are placed on HRF properties (e.g., smoothness). In the current context, a list of HRF morphological features (Table 1) are used to illustrate and compare model performance.

**Figure 2:**
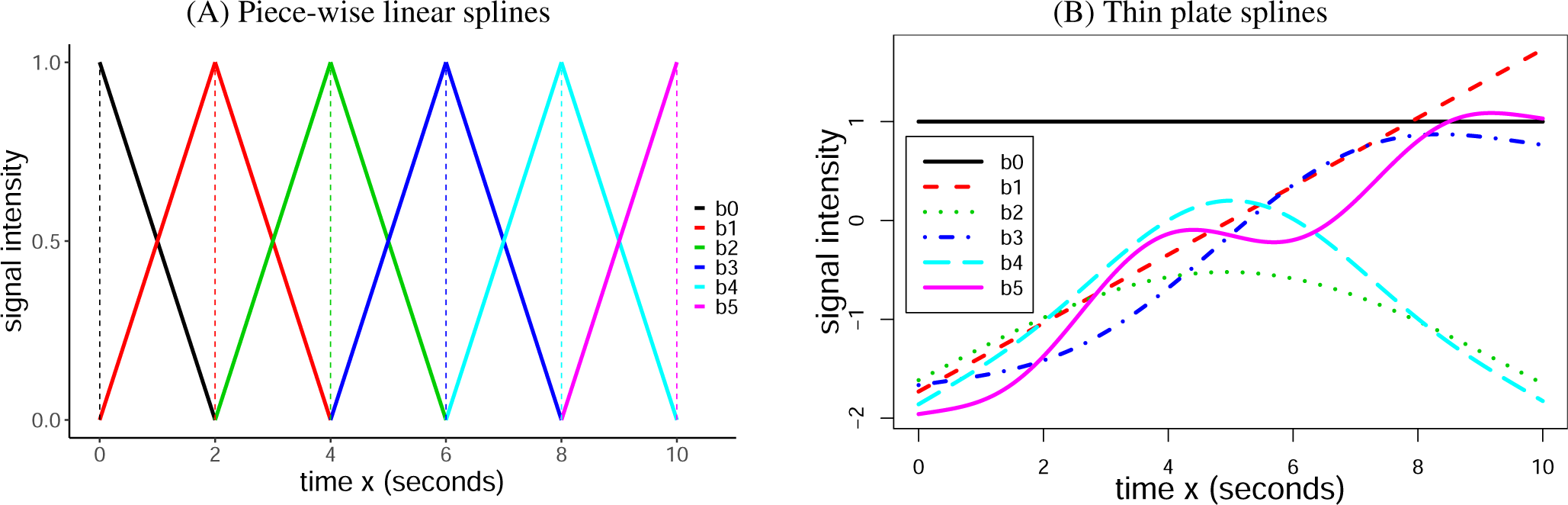
A set of splines *b*_0_(*x*)*, b*_1_(*x*)*, …, b_K_*(*x*) with *K* = 5. (A) Piece-wise linear spines in the form of tents (solid) or sticks (dashed). Each basis function characterizes a particular temporal interval centered on a knot except for the two end points. Although illustrated to cover an overall span of 10 s through a linear combination of these bases, the number of knots can be adjusted to accommodate any duration based on temporal resolution and model complexity. This type of basis function is used in neuroimaging software packages for individual-level modeling. (B) Thin plate splines. Unlike other smoothing splines (e.g., natural cubic splines), each of the thin plate splines spans the entire duration and is not tied to a specific knot; moreover, each thin plate incrementally adds more complexity. Intuitively, the first two bases, *b*_0_(*x*) and *b*_1_(*x*), capture the overall mean and linearity, respectively, while the rest catch the extent to which there is a nonlinear relationship.

**Figure 3:**
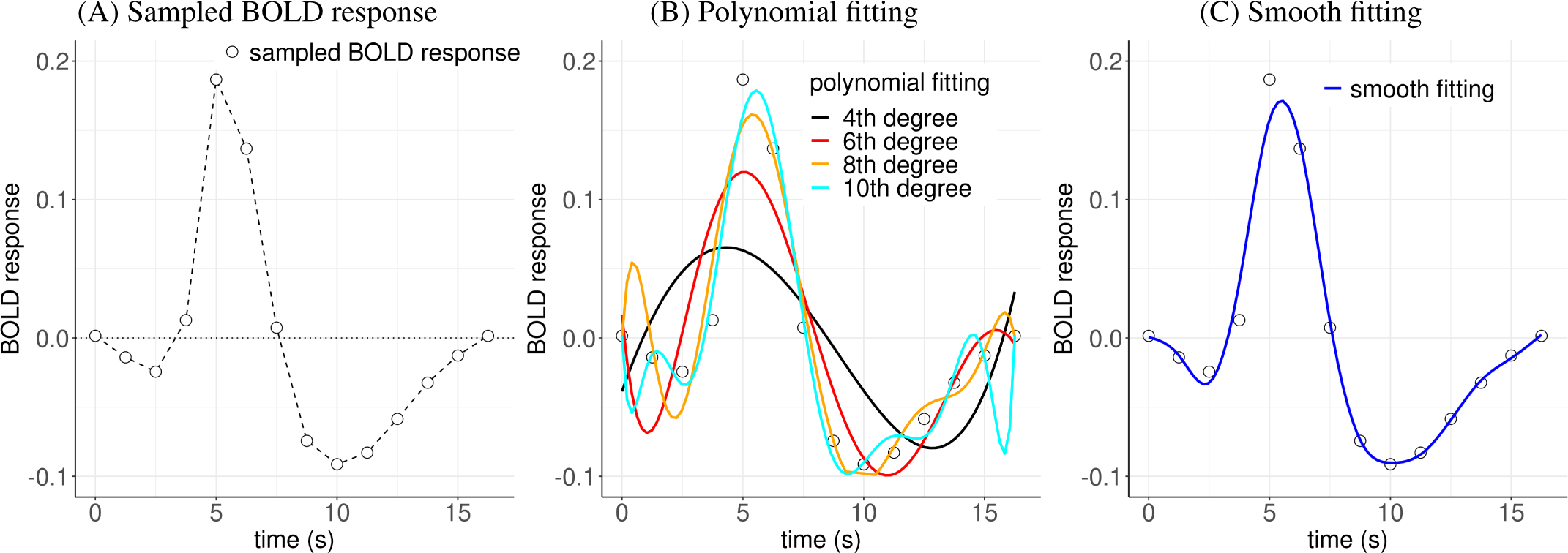
Comparisons of fitting BOLD response data. (A) A hemodynamic response (in units of percent signal change), sampled at 14 points (empty circles) with a time resolution of 1.25 s, was estimated from an experimental participant through a regression model. The sampled values can be viewed with linear piecewise interpolation (dashed line). These points have a “jagged” appearance, even though they are sampled from a presumably smooth HRF. (B) Fitting the data with polynomials introduces some smoothness but is also usually challenging: even though higher-order polynomials fit better to the original data, they may poorly make out-of-sample predictions and introduce extraneous features. (C) Modeling the HRF with smooth splines (e.g., thin plates) intends to achieve a counterbalance between fitting and predictive accuracy.

**Table 1:**
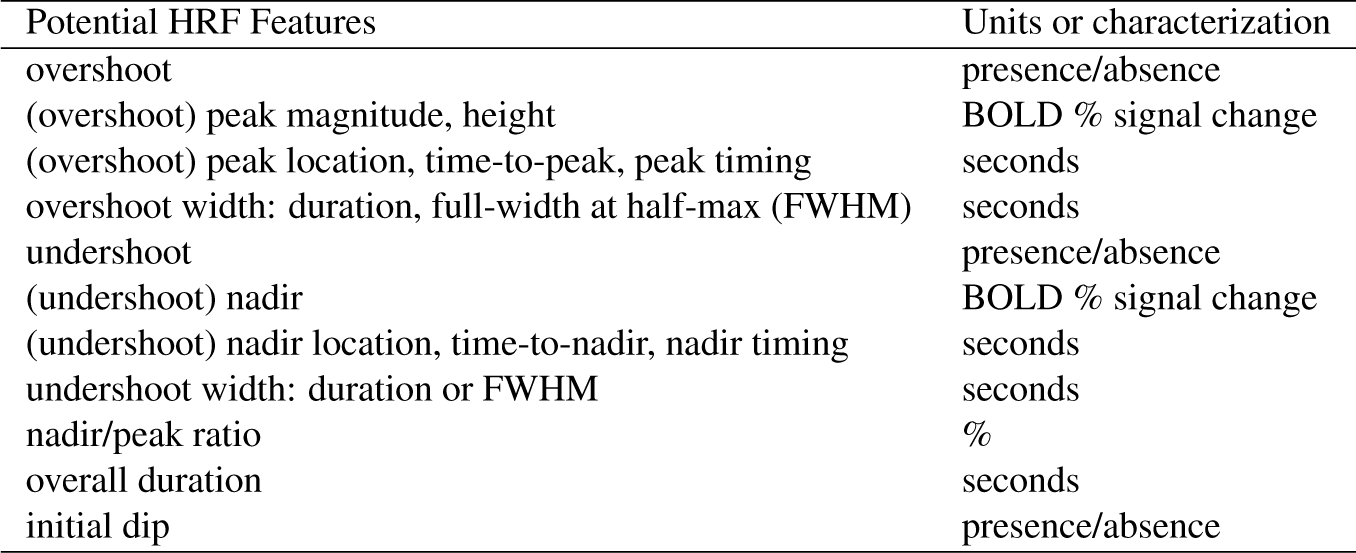
HRF morphological features considered.

| Potential HRF Features | Units or characterization |
| --- | --- |
| overshoot | presence/absence |
| (overshoot) peak magnitude, height | BOLD % signal change |
| (overshoot) peak location, time-to-peak, peak timing | seconds |
| overshoot width: duration, full-width at half-max (FWHM) | seconds |
| undershoot | presence/absence |
| (undershoot) nadir | BOLD % signal change |
| (undershoot) nadir location, time-to-nadir, nadir timing | seconds |
| undershoot width: duration or FWHM | seconds |
| nadir/peak ratio | % |
| overall duration | seconds |
| initial dip | presence/absence |

### 1.3 Preview: handling sampled HRFs at the population level

While HRFs are estimated with samples at the individual level, it is challenging to assess their morphology when brought into the population-level analysis. With a canonical HRF, the basic shape is predetermined. Thus, the focus at the individual level is usually on the regression coefficient as a scaling factor or a proxy for the peak magnitude, which is typically carried to the population level. When estimated through deconvolution with piecewise splines at the individual level, each HRF is represented by discrete samples (e.g., one at each TR). For instance, under the assumption of no BOLD response at the stimulus onset, with eight tents (Fig. 2) to cover a duration of 16 s (TR: 2 s), the estimated HRF from each individual is characterized by eight data points (i.e., regression coefficients). One could summarize each individual’s estimated HRFs via information reduction techniques, such as calculating the area under the curve at the overshoot, as an indicator of BOLD response (e.g., Beauchamp et al., 2003). However, such information reduction would result in information loss, negating the original goal of capturing the whole HRF profile. Alternatively, HRF samples at time points can be brought to the population level as a repeated-measures factor (Chen et al., 2015) to be analyzed under a conventional AN(C)OVA or linear mixed-effects modeling framework, as implemented in the programs 3dMVM (Chen et al., 2014) and 3dLMEr (Chen et al., 2013) in AFNI. Then, an interaction approach is adopted to examine statistical evidence: by treating the HRF samples as the levels of a repeated-measures factor, the presence of the BOLD response can be assessed through the main effect (one-way interaction), and HRF differences across groups and conditions can be inferred through two-way interactions with the repeated-measures factor. It has been shown that detection through this interaction approach achieves substantially higher sensitivity than the canonical approach (Chen et al., 2015).

While the above approach of comparing sampled HRFs has benefits, it may be possible to improve on these approaches to reducing information loss and also perhaps increasing predictive accuracy. For example, neuron action potential with which the sampled HRF is associated can be approximated through a smooth curve modeled by differential equations (Hodgkin and Huxley, 1952). However, discretely sampled HRFs ignore the smoothness aspect, which may make the response modeling susceptible to noise. In other words, the sampled HRF model is fit to the raw individual-level data without regularization through an interaction within an AN(C)OVA (Chen et al., 2015), even if that results in jagged appearances (Fig. 3A) and over-fitting of the estimated HRF. Statistical inferences are then based on those sampled time points without their sequential information taken into consideration. For instance, the serial order among three consecutive time points of *k −* 1, *k*, and *k* + 1 TRs (*k* = 1, 2*, …*) are not preserved since they are simply treated as three levels of a repeated-measures factor in the AN(C)OVA model. Hence, the interaction approach is disconnected from the underlying BOLD mechanism, which may cause the loss of sensitivity and inferential efficiency. Here, we adopt a smooth modeling approach: let the data determine the BOLD response morphology for each voxel and subsequently apply a regularization step. Specifically, we extend the individual-level HRF estimation method through deconvolution with splines (either piecewise linear, as in Fig. 2A, or cubic splines) to the population level. Moreover, with minimal assumptions that play the role of adaptive regularization, we impose a constraint on the HRF smoothness that should reduce the influence of noise and prevent the informed profile from over-fitting. In other words, instead of assessing how consistent an HRF is with data alone, we assess how consistent the HRF is with the data as well as its smoothness.

Previous work on HRF estimation has explored data-driven approaches or a smoothness constraint using, for example, a Gaussian process prior (Goutte et al., 2000; Ciuciu et al., 2003; Eickenberg et al., 2017), cubic smoothing splines (Zhang et al., 2007), B-splines (Degras and Lindquist, 2014), a canonical HRF combined with its temporal derivative (Elbau et al., 2018), wavelet bases (Van De Ville et al., 2004; Khalidov et al., 2011), a biophysically informed HRF (Rosa et al., 2015), Tikhonov regularization (Zhang et al., 2007; Casanova et al., 2008; Casanova et al. 2009; Zhang et al. 2012), spatial regularization (Badillo et al., 2013; Chaari et al., 2013; Zhang et al., 2018), cross validation (Zhang et al., 2013), and nonlinear optimization (Pedregosa et al., 2015). Some of these methods were applied to individual-level modeling for task-based experiments (Goutte et al., 2000; Ciuciu et al., 2003; Zhang et al., 2007; Chaari et al., 2013; Pedregosa et al., 2015) and for resting state data (Wu et al., 2021); Cherkaoui et al., 2021). Other methods have been adopted at the region level (Chaari et al., 2013; Zhang et al., 2012; Badillo et al., 2013; Zhang et al., 2013; Zhang et al., 2018) or developed for information extraction through information reduction of HRF to two or three morphological features (Zhang et al., 2012; Zhang et al., 2013).

Our current work extends these previous methodologies, but differs in two key aspects. First, at the individual level, we directly estimate the HRFs using deconvolution with piecewise splines without imposing smoothness or any other regularization. Second, we carry the estimated HRFs to the population level where we apply regularization. In doing so, we intend to improve population-level inferences for cross-group and cross-condition comparisons based on the full HRF, not on one or a few morphological features.

In addition to laying out the modeling framework of the smooth HRF, we will address the following questions using a dataset of participants performing a sustained attention task during FMRI scanning:

1. How much does the HRF shape vary across regions, tasks, conditions, and groups?
2. Does the smooth HRF approach improve effect detection sensitivity and efficiency?
3. How useful is the HRF shape information in verifying the presence of an effect?
4. Is the whole brain involved during a simple task? A previous “deep data” study by Gonzalez-Castillo et al. (2012) indicated that most of the brain was involved in a simple block experiment of visual stimulation at the individual level. Compared to such an experiment with many blocks (e.g., 500), in which the average BOLD response is relatively strong and reliable, an event-related experiment can be challenging to model due to the much weaker signal. Does the characterization of the HRF shape provide more sensitivity and possibly reveal similarly widespread fluctuations in most brain regions at the *population* level in a rapid event-related design?
5. Does the HRF estimation approach provide evidence for BOLD response in white matter? Addressing this is particularly important, as average white matter signals are often used to construct confound regressors for motion and other physiological effects in resting state FMRI.

We will first discuss the smooth HRF approach with the specific goal of assessing the hemodynamic response at the population level. Next, we demonstrate the analytical strength of this approach using an experimental, task-based FMRI dataset. Finally, we will summarize the three methodologies of canonical, sampled, and smooth HRF, and compare their pros and cons.

## 2 Estimating population-level HRF through smoothing splines

In this section, we describe the mathematical background and formulation of the smooth HRF modeling approach; the next section describes its application. We start with a simple case of an FMRI study with one task condition and one group of participants^2^. Let {(*t_k_, y_sk_*): *s* = 1, 2*, …, S*; *k* = 0, 1*, …, K*} be a set of HRF samples under one condition from *S* individuals (indexed by *s*), which are estimated at *K* + 1 knots (indexed by *k*) from an individual-level regression analysis. Specifically, *y_sk_* is the regression coefficient for the HRF sample of *s*th individual at *k*th time point. Then, (*y_s_*_0_, *y_s_*_1_, *…, y_SK_*) are the sampled data that characterize the *s*th individual’s HRF profile, while (*t*_0_, *t*_1_, *…, t_K_*) are the corresponding knot locations (or times, usually taking the incremental values in the unit of TR counts or seconds). For example, *K* = 13 would correspond to having 14 knots that span 13 TRs for a condition, and *t_k_* = *k, k* = 0, 1*, …,* 13. As noted above, the HRF estimation occurs independently at each voxel in the dataset.

### 2.1 Model formulation

We assume that the BOLD response *y* for a condition can be characterized as an (initially) unknown, continuous and *smooth* function *y* = *h*(*t*) of time *t*, where the function *h*(*t*) has up to second-order derivatives (defining its minimal smoothness). Visually, a smooth function can be described as a curve without sharp bends, or mathematically defined as at least first order differentiable. Here, we make the practical assumption of having second-order differentiability partly for mathematical and algorithmic convenience and partly for its visual interpretability (e.g., smoothly connecting discrete points with only moderate smoothing). Regarding common splines currently used at the individual level in the field, piecewise linear splines such as the tent/stick basis set in AFNI and FIR basis in FSL/SPM do not meet this smoothness requirement because they have sharp corners (i.e., a discontinuity in their first derivative) at the knots (Fig. 2A). However, the cubic splines available in AFNI do satisfy the smoothness criterion. We note that the notations of *t_k_* and *y_k_*, introduced earlier with the subscript *k* added, are meant to associate the discrete (sampled) quantities with their continuous counterparts of *t* and *y*, respectively, at the *k*th knot. Here and below, when *y* appears with a single subscript *k*, we have removed the index *s*, letting *y_k_* refer to the response at the *k*th knot for an unspecified individual.

We start by illustrating the method with a single generic individual. The model is constructed for the hemodynamic response sample dataset {(*t_k_, y_k_*)} as follows:

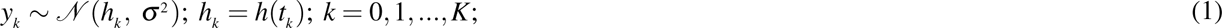

where *h*(*t*) is the theoretical HRF, *h_k_* is its discrete time-sampled value at the knot *t_k_*, and *σ* is the standard deviation for the sampled data *y_k_*. As a concrete example, Fig. 3A illustrates a set of data *y_k_* sampled at time points (in TR) *t_k_* = *k, k* = 0, 1*, …,* 13, and Fig. 3B-C show possible ways to estimate an HRF *h*(*t*).

Distinct from curve fitting, “smooth fitting” through regularization is an adaptive calibration process for interpolation (Fig. 3C). Due to limited samples and noisy fluctuations, the original BOLD response data *y_k_* tend to have a rough appearance (Fig. 3A). When confined to those samples, statistical inference cannot be extended to the intervals beyond those sampled time points. For example, the predictive accuracy beyond the sample data points is important in estimating the peak height due to the high sensitivity to its location. The traditional nonlinear smooth fitting through polynomials is also daunting (Fig. 3B): order selection for polynomials can be complicated and arbitrary, and the fitted curve at a particular point may be sensitive to both small, local fluctuations and distance measures (e.g., Runge’s phenomenon). Instead of fully relying on the data and tracing the trajectory exactly through each data point (Fig. 3A), we seek to fit an approximate curve using a regularized process (Fig. 3C): use the data *y_k_* to inform and learn the underlying HRF *h*(*t*) morphology while at the same time containing the risk of over-fitting and the vulnerability to sampling uncertainty through imposing a neurologically reasonable constraint of smooth transitions. Moreover, because of the assumption of HRF smoothness, we avoid taking the risk of two extremes: over-fitting (fully trusting HRF samples or high-order polynomial fitting) and under-fitting (e.g., canonical HRF, linear or low-order polynomial fitting). As a result, we expect to achieve a high predictive accuracy and to improve statistical inferences over the entirety of each HRF profile as opposed to being restricted to the sampled time points *t_k_*.

### 2.2 Fitting the model with smooth splines

To achieve smooth fitting, we adopt a modeling strategy at the population level through splines. Many types of smooth splines are available, differing in the expression of their basis functions while mathematically equivalent. Here, we focus on utilizing thin plate splines (Fig. 2B), given their numerical flexibility and performance (Wood, 2003). Additionally, thin plate splines are organized in terms of increasingly nonlinear complexity across spline order, which is convenient for approximation and interpretability. Estimation starts with as many splines as the number of knots, *K*. To reduce computation and memory overhead, eigen-decomposition is adopted with *K* + 1 splines. The number of splines is then reduced to the most *P* + 1 important components to preserve much of the nonlinearity among the conventional thin plate bases, but at considerably greater computational efficiency for large datasets. Unlike other spline sets, thin plate splines are not tied to a set of knots (Fig. 2B). Because of the component reduction process, the number of adopted thin plates is usually less than the number of knots, *K*, adopted at the individual level (where *K* is determined by FMRI sampling rate and assumed total HRF duration). In other words, we will often have *P ≤ K* (such as *P* = 9 with *K* = 13). Another unique (and convenient) feature of thin plate splines is the structure of their first two bases: overall mean (solid black, Fig. 2B) and linear (dashed red, Fig. 2B) splines correspond to baseline and linearity. We now express the smooth HRF *h*(*t*) as a linear combination of *P* + 1 thin plates within the time interval *t ∈* [*t*_0_, *t_K_*],

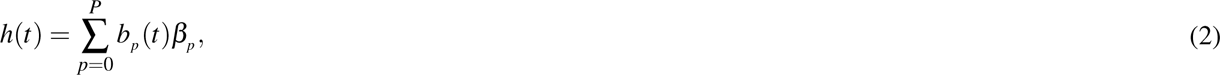

where {*b_p_* (*t*): *p* = 0, 1*, …, P*} are *P* + 1 thin plates (indexed by *p*). The model (1) can then be reformulated,

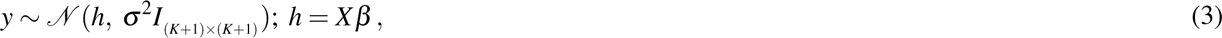

where *y* = (*y_0_, y_1_, …, y_K_*)′, *h* = (*h*(*t*_0_)*, h*(*t*_1_)*, …, h*(*t_K_*))′, *β* = (*β*_0_, *β*_1_, …, *β_P_*)′, and *X* is a (*K* + 1) *×* (*P* + 1) matrix whose (*k, p*)th element is composed of *b_p_* (*t_k_*). By replacing the unknown but presumably smooth nonlinear function *h*(*t*) in (1) with the basis expansion (2), we have successfully converted a nonlinear relationship (1) into a linear formulation (3) with the model matrix *X* composed of the basis functions evaluated at the time points *t_k_*.

A regularization process is adopted to balance between nonlinearity complexity and roughness. Spline fitting has a long history of applications in many fields, including individual-level modeling in neuroimaging and signal processing in general. However, the adoption of smooth fitting via regularization is relatively recent in statistical modeling (Hastie and Tibshirani, 1990; Wahba, 1990; Wood, 2017). At one extreme, fitting with a high number of splines could capture more complex associations and lead to many rapid turns with large curvature, but it may then over-fit the data and capture noise. For example, with *K* + 1 time points, we could adopt *K* + 1 splines and fit with *K*th order polynomials. Such a fit would likely suffer from poor predictive accuracy when applied to out-of-sample data (Fig. 3B). At the other extreme, too few splines could under-fit nonlinear associations. For instance, with the first two thin plates—*b*_0_(*t*) = 1 and *b*_1_(*t*) = *t*, Fig. 2B—one could fit any data in a traditional linear regression model through a straight line (infinitely smooth) with no turns (i.e., zero curvature). However, such a fit would not be a compelling choice especially for HRFs.

The balance between nonlinearity and roughness can be achieved through regularization. As the second derivative of a function is associated with its curvature or concavity, a positive second derivative corresponds to an upwardly concave, while a negative second derivative represents downward concavity. Thus, we control the smoothness or curvature of the HRF *h*(*t*) by tuning the following penalty term on the integrated square of its second derivative (Wood, 2017; Chen et al., 2021a),

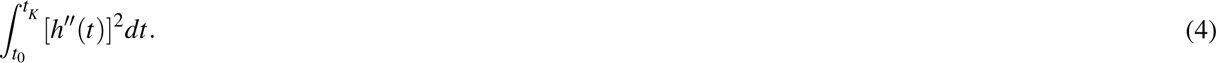

The metric (4) is considered a natural measure of function roughness or wiggliness (Wood, 2017), which can be expressed as a regularization process among all the spline weights excluding the first two (corresponding to baseline and linear trend, whose second derivatives are zero): (*β*_2_, *β*_3_, *…, β_P_*). An example of the impact of this regularization on the original data (Fig. 3A) is illustrated in Fig. 3C. One trivial case is the following: if only the first two splines of *b*_0_ (*t*) = 1 and *b*_1_ (*t*) = *t* are adopted, the smooth fitting model (3) degenerates into a linear regression: As *h^′′^*(*t*) = 0 in this case, the penalty (4) becomes zero and no regularization is applied.

Conceptually, the regularization in smooth fitting can be viewed in three ways that provide insight into its modeling applications. First, as the first two splines among the thin plates are overall mean and linearity (Fig. 2B), the penalty (4) on *β* is essentially an *L*^2^ regularization imposed on those spline weights other than the first two elements, *β*_0_ and *β*_1_ (Wood, 2017; Chen et al., 2021). That is, the information is regularized across the nonlinear elements (*β*_2_, *β*_3_, *…, β_P_*)′. Algorithmically, the penalty (4) can be applied with a tuning or smoothing parameter through partial least squares. Thus, the HRF *h*(*t*) can be expressed with the *β* weights in (2) and then numerically estimated through a process similar to the conventional ridge regression. Second, the penalty (4) can also be formulated as a linear mixed-effects (LME) model with the spline weights (*β*_2_, *β*_3_, …, *β_P_*)′ as random effects. In this case, we can readily adopt the numerical schemes from the traditional LME modeling approach. Lastly, under the Bayesian framework, the penalty (4) is formulated as a prior assumption for the spline weights (*β*_2_, *β*_3_, …, *β_P_*)′. All these three frameworks are practically useful in effect estimation and statistical inference. In the end, the regularization is implemented through imposing a Gaussian prior on those higher order splines, which intends to strike a balance between over- and under-fitting. For technical details, refer to Wood (2017).

A few features differentiate the HRF model (2), which is fitted with smooth splines, from the typical HRF estimation methods at the individual level as currently implemented in neuroimaging packages. First, most basis sets (e.g., stick/tent in AFNI, finite impulse response in FSL/SPM) are not smooth splines. Second, each of the thin plate splines covers the entire interval, and the degree of modeled nonlinearity increases with the number of bases. In contrast, each of the basis sets adopted in typical individual-level modeling is piecewise in the sense that each basis is local to one knot sharing the same functional form (e.g., linear or cubic polynomial) rather than increasing in complexity. Third, the smoothness of the HRFs fitted in the model (2) is further regularized through the roughness measure (4). In contrast, no regularization is applied in the conventional HRF estimation approach through splines. For example, the cubic splines adopted in AFNI, even though smooth, are used to generate regressors with no subsequent regularization in the modeling process.

### 2.3 Model extensions

Now, we refocus our attention on estimating a smooth HRF *h*(*t*) at the population level. With the individual subscript *s* re-included, a hierarchical model is constructed at the population level with the set of individual-level HRFs {(*t_k_, y_sk_*)} as input:

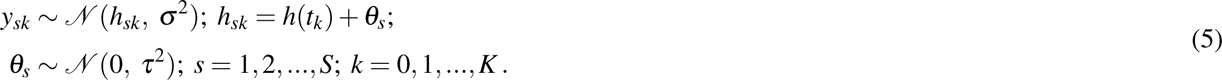

Here, *h_sk_* is the value of HRF *h*(*t*) associated with the *s*th individual at the knot *t_k_*, *σ* ^2^ is the intra-individual variance. *θ_s_* codifies the individual-specific intercept, and *τ*^2^ represents the inter-individual variance. The model platform (5) can be extended to include categorical variables, such as multiple groups of participants (e.g., controls versus patients), where the investigator may be interested in comparing HRFs between groups or simply accounting for group differences in HRFs. Similarly, quantitative variables (e.g., age) can also be incorporated.

Statistical evidence for the predicted smooth HRFs is inferred as follows. A *χ*^2^-statistic is adopted to assess the evidence for each estimated HRF using its uncertainty within a Bayesian framework (Wood, 2013; Wood, 2017), where the null hypothesis posits that the HRF is a horizontal line: *h′* (*t*) = 0. When comparing two HRFs, the expected responses at any time *t* are estimated per the model (2). The uncertainty band of each estimated HRF is expressed as the standard error at every time point.

### 2.4 Modeling implementation for whole-brain voxel-wise analysis

The program 3dMSS, which implements the smooth HRF modeling approach for whole-brain voxel-wise data analysis, is publicly available in AFNI. It uses the R (R Core Team, 2022) package gamm4 (Wood and Scheipl, 2020), with thin plate splines as a default. Other basis functions such as cardinal cubic and tensor product splines are also available. The data analysis is executed through a shell scripting interface. All explanatory variables and input data (in text format for 1D and 2D files or NIfTI format for 3D neuroimaging data) are prepared in a table in the R long data format. Run-time may range from minutes to hours, depending on the amount of data, spatial resolution, the number of predictors, model complexity, and computational capacity.

We make the following recommendation for choosing the number of basis functions. At the individual level, one should achieve the highest possible time resolution for the estimated HRFs. For example, one may use deconvolution with 8 piecewise splines to cover a response duration of 16 s for a scanning TR of 2 s. At the population level, when 10 or more samples are available for estimated HRFs, we recommend 10 smooth splines (the default for 3dMSS), as a suitable choice to capture all major BOLD response shape features. With less than 10 HRF samples, choose the number of bases to be close or equal to the number of HRF samples. Thanks to the eigen-decomposition process, this recommendation keeps a balance between adequate smooth fitting and unnecessary computational cost.

HRF predictions and statistical inferences are provided in the 3dMSS output. The statistical evidence for each HRF or for the comparison between two HRFs, is assessed through a *χ*^2^-statistic based on all basis weights associated with the nonlinear splines, (*β*_2_, *β*_3_, *…, β_P_*) in the model (2), and their variance-covariance matrix (Wood, 2017). As the effective degrees of freedom vary across the spatial units (due to eigen-decomposition-based component reduction performed at the voxel level), each *χ*^2^-statistic is transformed to have 2 degrees of freedom for convenient result storage. To make predictions, the user can specify the time resolution for visualization and other variables in a long data format and obtain both the estimated smooth HRFs and their standard errors in the output. One can then interactively examine the HRF morphology and comparisons at the voxel level through, for example, the AFNI graphical user interface or other tools.

To briefly summarize, we assess three modeling approaches (Table 2): 1) the canonical HRF, which has fixed shape and scalar comparison; 2) the sampled HRF, which has a flexible shape and ANOVA-based comparison; and 3) the smooth HRF, which has a flexible shape, fit-regularization and curve-based comparison. We hypothesize that the canonical approach will be the least accurate due to under-fitting, followed by sampled and smooth HRF approaches. One may expect that the sampled HRF approach could have similar performance to its smooth counterpart, but the smooth HRF approach should have two advantages. First, HRF samples tend to contain some amount of noise because of their dependence on sample sizes and model quality. In contrast, as the bias–variance trade-off shows, the regularization in smooth HRFs should reduce noise sensitivity, and yield higher out-of-sample predictive accuracy and reproducibility. Second, and perhaps more important, the regularization should also increase inferential efficiency, facilitating the ability to make comparisons.

**Table 2:** Comparisons among different HRF modeling assumptions.

| Canonical HRF | Sampled HRF | Smooth HRF |
| --- | --- | --- |
| fixed features: total duration, time-to-peak, overshoot width, time-to-nadir, undershoot width, peak magnitude, peak/nadir ratio, curvature | fixed total duration | fixed total duration, assumed smoothness |

## 3 HRF estimation with an experimental dataset

We now assess the smooth HRF modeling approach in comparison with the other two established methods of canonical and sampled HRFs. The adopted dataset has been analyzed in a prior study (Pagliaccio et al., 2017a) of a global-local selective attention task during FMRI scanning (Weissman et al., 2006). Briefly, 87 participants (mean age: 18.3, SD: 4.2, range: 8-25 years, 58% female) passed quality control. Of these, 43 were either diagnosed with bxipolar disorder or at familial risk for bipolar disorder (BP), and 44 were healthy volunteers (HV). On each trial, participants saw a large letter (H or S) made up of smaller letters (Hs or Ss) and were asked to identify, by button press, the large “global” letter or the small “local” letters, depending on the run-level instructions. In half of the trials, the displays were congruent (e.g., a large H consisted of small Hs), while in the other half of the trials, the displays were incongruent (e.g., a large S consisted of small Hs). Stimulus ordering was pseudorandomized. Each stimulus appeared for 200 ms, followed by a 2300 ms fixation/response window, and an additional inter-stimulus interval jittered in TR increments (1250 ms) with a decay curve. The total trial length ranged from 2.5 to 16.25 s (mean *≈* 4.0 s). Six scan runs (304 time points each) were acquired per participant, each composed of 92 trials (46 congruent and 46 incongruent), alternating the task goal of identifying global or local letters. The scanning sequence used a TR of 1250 ms with a voxel size of 3.75 *×* 3.75 *×* 5.00 mm^3^. Detailed information on data acquisition and sequence parameters can be found in Pagliaccio et al. (2017a).

### 3.1 Data preprocessing and individual-level analysis

Data were analyzed using AFNI (Cox, 1996; version 22.1.14) with standard preprocessing via afni_proc.py. Processing commands and scripts are available online (https://github.com/afni/apaper_hrf_profiles). First, both skullstripping each individual’s anatomical image and the estimation of its nonlinear alignment to the MNI 2009c template were performed using 3dQwarp through @SSwarper (Cox and Glen, 2013). The initial processing steps included removal of the first 6 TRs, as well as despiking and slice-timing correction. EPI-anatomical alignment was performed using the lpc+ZZ cost function (Saad et al., 2009), while also applying EPI brightness unifizing to aid feature matching, and checking for left-right flips (Glen et al., 2020). To address head motion, EPI volume registration was performed by rigid alignment to the “MIN_OUTLIER” reference base (the volume with fewest outliers), and the six head motion series were saved to use in individual-level regression analysis. All alignment steps were concatenated into a single transform before applying to minimize smoothing, and the final EPI voxel dimension was set at 3.5 mm isotropic. A whole-brain mask was created as the intersection of the EPI and the anatomical and was applied in subsequent steps (Taylor et al., 2018b). The EPI time series were blurred within the brain mask to FWHM = 6.5 mm, and then were scaled to local average for interpreting effect estimates as percent signal change (Chen et al., 2017).

Individual-level regression models were fitted through generalized least squares with a temporal autocorrelation structure of ARMA(1,1) with 3dREMLfit. Baseline, low-frequent drifts and motion regressors were included per run. To further reduce motion effects, censoring was carried out to exclude time points where the Euclidean norm of the first difference in motion effects (Enorm) was > 0.3 mm, or where the fraction of outliers within the brain mask at a particular time point was > 5%. Sessions were excluded if greater than 15% of TRs were censored for motion/outliers or if the individual’s behavioral performance was below 70% accuracy. Additional quality control was conducted using afni_proc.py’s QC HTML (Reynolds et al., 2023). This included visual assessment of the EPI FOV coverage, of volumetric alignment (EPI-anatomical and anatomical-template), and of the full *F*-statistic map of the overall regression model. Grayplots of residuals and volumetric radial correlation plots were examined for FMRI signal variance and local correlation patterns that could be indicative of high motion/scanner coil artifacts, respectively. Individual and combined stimulus plots and the regression matrix were examined to confirm the accuracy of timing files and appropriately spaced stimulus events. All population-level analyses were restricted to a whole-brain mask where at least 90% of participants had usable data (33,495 voxels).

Two established modeling approaches (canonical and sampled HRFs) were adopted at the individual level. The first one used the canonical HRF with regressors of interest (congruent, incongruent) created through a gamma variate. The second one estimated HRFs through deconvolution with piecewise linear splines (tent function); that is, each condition was modeled with 14 basis functions that covered a sampled HRF with a duration of 16.25 s (13 TRs) window starting 2.5 s before stimulus onset. The first two TRs were used to capture pre-stimulus dips in BOLD response that have been associated with attention lapses (Weissman et al. (2006).

### 3.2 Modeling at the population level

The input data at the population level were structured as follows: For the canonical approach, an effect estimate (regression coefficient) was available for each of the two conditions and each of the 87 participants, resulting in 174 total 3D volumes. For the sampled and smooth HRF approaches, each participant had 14 effect estimates as a sampled HRF per condition that spanned over 13 TRs, leading to 2,436 total 3D volumes. Sex and age were included as covariates in all cases.

Three modeling approaches were adopted at the population level. First, for the canonical method, an ANCOVA was formulated to examine canonical HRF peak magnitude using the AFNI program 3dMVM with three factors (group, condition, and sex) and one quantitative covariate (age). Second, for the sampled HRF method, a similar ANCOVA model with 3dMVM was performed but with an extra factor that had 14 levels for the HRF-sampled time points. Each effect of interest (e.g., group/condition difference) was assessed by the interaction of the relevant variables (e.g., group, condition) with the HRF factor (Chen et al., 2015). Third, for the smooth spline method, the HRF for each group under each condition was fitted at the 14 temporal samples from the individual level with sex and age as covariates using the program 3dMSS. Scripts are available at https://github.com/afni/apaper_hrf_profiles.

### 3.3 Comparisons among the three modeling approaches

Here, we adopt a “highlight-but-not-hide” approach when visualizing results, to utilize available information more fully, to be less sensitive to the relatively arbitrary threshold, and to provide a more complete description of the results (Taylor et al., 2022b; Chen et al., 2022b). In the highlighting approach, transparent threhsolding is implemented: suprathreshold results are opaque while subthreshold ones are still shown, but with increasing transparency as statistical strength weakens. To explore statistical evidence broadly, the voxel-level threshold in displayed images was set to *p* = 0.01. For direct visual comparison across the three modeling approaches, no spatial clustering was applied. Displayed colors indicate the strength of statistical evidence. The predicted HRFs using smooth splines were sampled at quarters of the TR (i.e., a time resolution of 0.3125 s) for visualization. In each image below, the neurological convention was adopted: the left side of the brain is shown on the left of the image. Additionally, RAI-DICOM conventions are used for reporting coordinates: a negative *X* denotes the right hemisphere; negative *Y*, the anterior; and negative *Z*, the inferior.

The omnibus results emerge in the expected regions for the group and condition effects (Fig. 4). For example, the motor and medial prefrontal cortex regions reveal group differences, and the bilateral dorso-lateral PFC and parietal regions show condition effects. Previous work using canonical and sampled HRFs did not identify group-by-condition interactions (Pagliaccio et al., 2017a; Pagliaccio et al., 2017b), which may have possibly been due to the lack of sensitivity of the established approaches.

**Figure 4:**
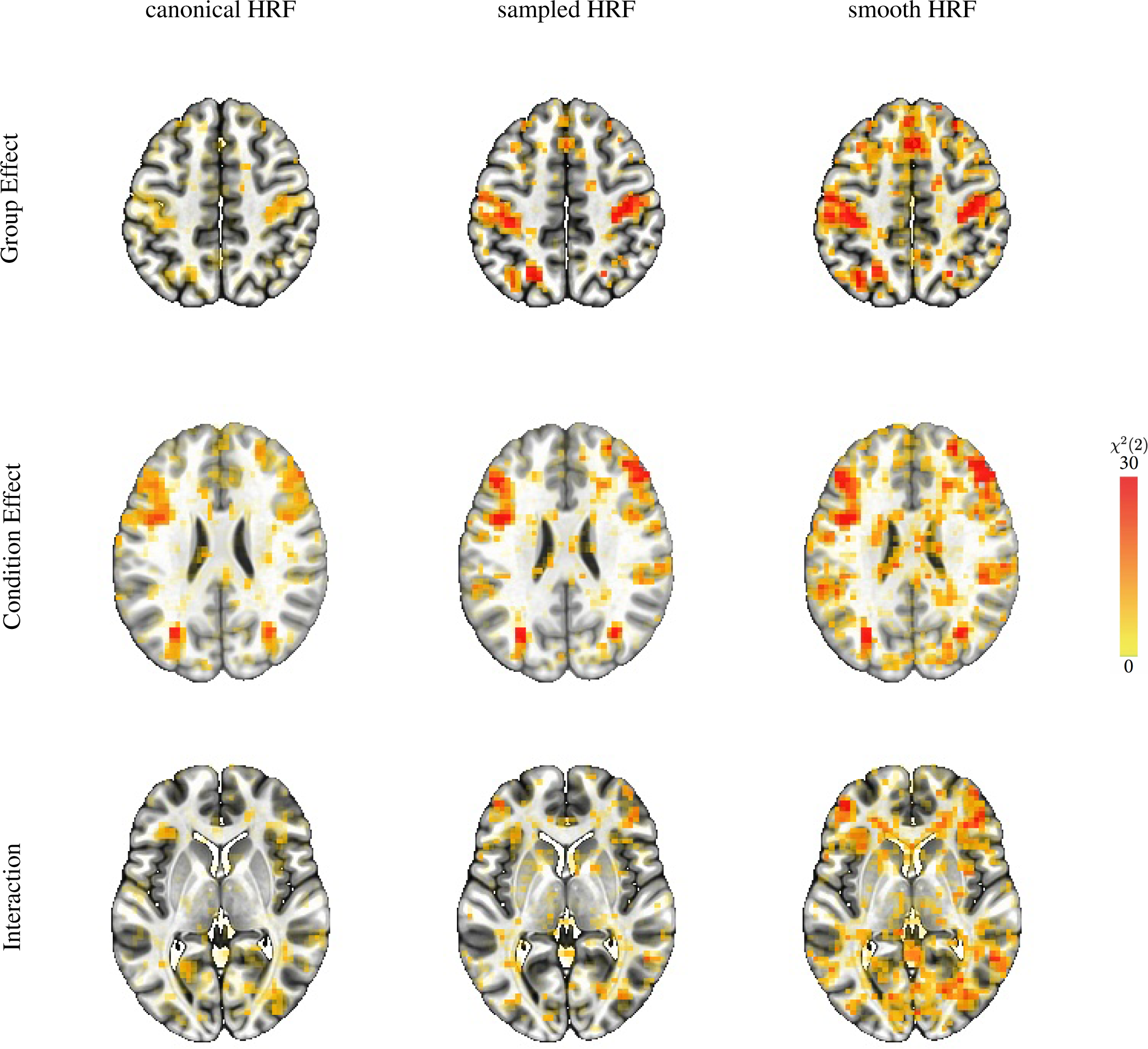
Statistical comparison of main effects and interactions among the three modeling approaches. Overlay coloration shows *χ*^2^(2) values; transparent thresholding at *p* = 0.01 here and below corresponds to *χ*^2^(2) = 9.2. The three different axial slices (rows from the top, downwards) at *Z* = 49, 25, and 4 are selected to illustrate the statistical evidence comparisons across the modeling approaches (columns) for, respectively, differences between groups and conditions, and the group-by-condition interaction.

These overall results confirm our prediction about the sequential order of the model performance. Fig. 4 shows the *χ*^2^ results for the group, condition, and interaction effects for each HRF modeling approach. The superior performance of the sampled HRF over the canonical HRF is consistent with our previous investigation (Chen et al., 2015). Noticeably, the smooth HRF approach was statistically more sensitive than the sampled HRF for the “omnibus” evaluation of the two main effects and the interaction. The improved sensitivity is especially beneficial for higher-order effects such as interactions (third row, Fig. 4) than main effects (first two rows, Fig. 4) because the former generally require much larger sample sizes than the latter (Leon and Heo, 2009; Gelman et al., 2020). As is common in result reporting in the field, Fig. 4 only shows one aspect of information: statistical evidence. However, it is equally – if not more – important to display the associated effects (Chen et al., 2017; Chen et al., 2022b). As illustrated below, the HRF estimation approaches make powerful visualization available and allow researchers to evaluate information beyond statistical evidence: Figs. 5-8 display the full HRF curves at a voxel, and Fig. 9 shows the voxelwise features across the brain.

**Figure 5:**
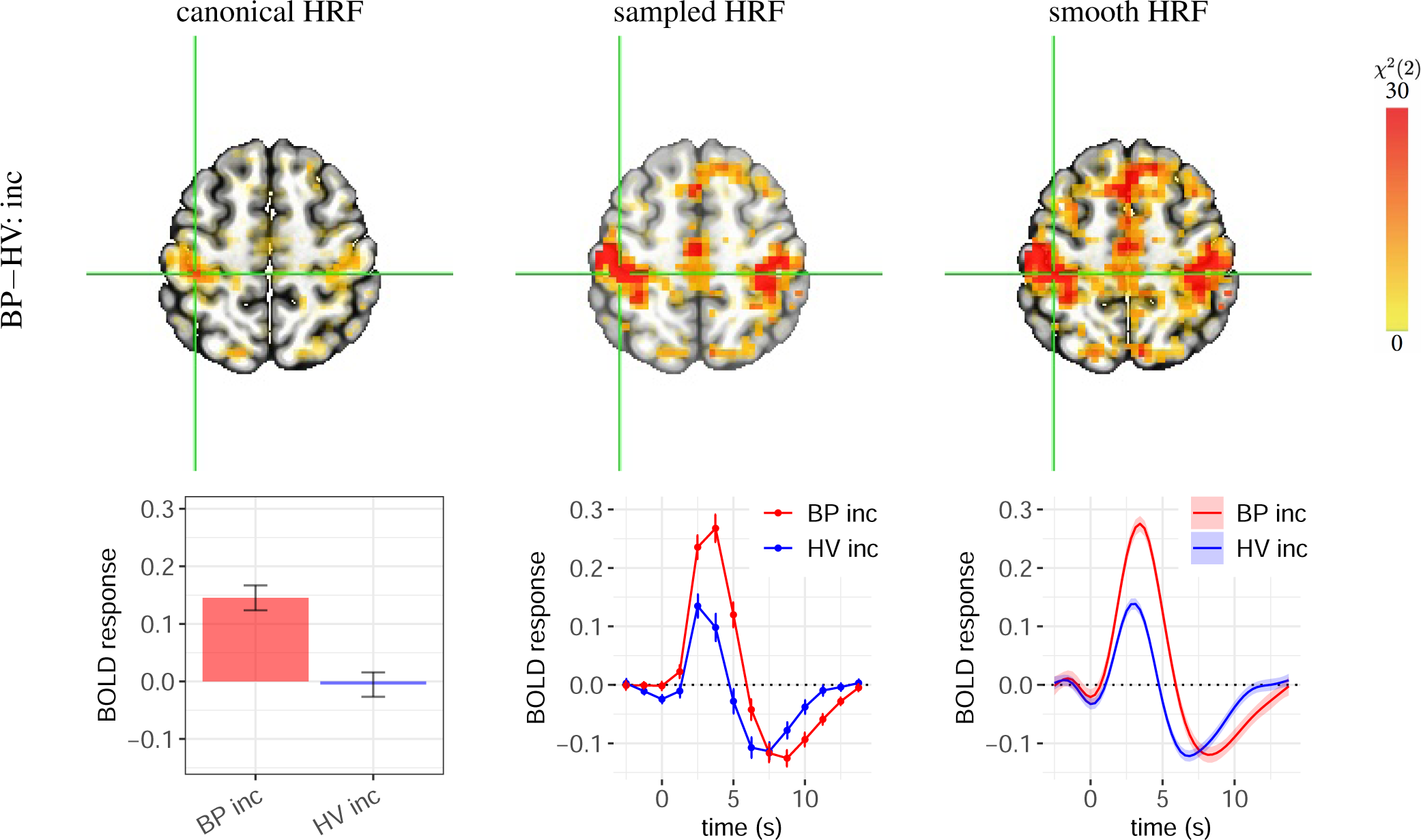
Case A: roughly compatible statistical evidence, but with important differences in effect estimates. Group differences are shown for the incongruent condition at (*X,Y, Z*) = (39, 29, 56) in the left postcentral gyrus. (top row) All three approaches showed strong statistical evidence for the presence of a group difference. (bottom row) The effect estimation of group difference differs importantly across methods. In each panel, vertical bars and shaded bands indicate standard error. As the estimated HRFs are largely similar to the canonical version, the estimated group contrast (0.15 *±* 0.03) from canonical HRF (first column) is roughly compatible with the HRF peak differences from the sampled and smooth HRF estimates. However, as the canonical HRF assumes a peak at 5 s, slightly later than what was revealed through HRF estimation, it substantially underestimated effects for both groups. Anticipation effects occurred about one TR before stimulus onset as illustrated by the HRF estimation approaches. HRFs for the other condition (BP con, and HV con) and neighboring voxels shared similar profiles (not shown here).

**Figure 6:**
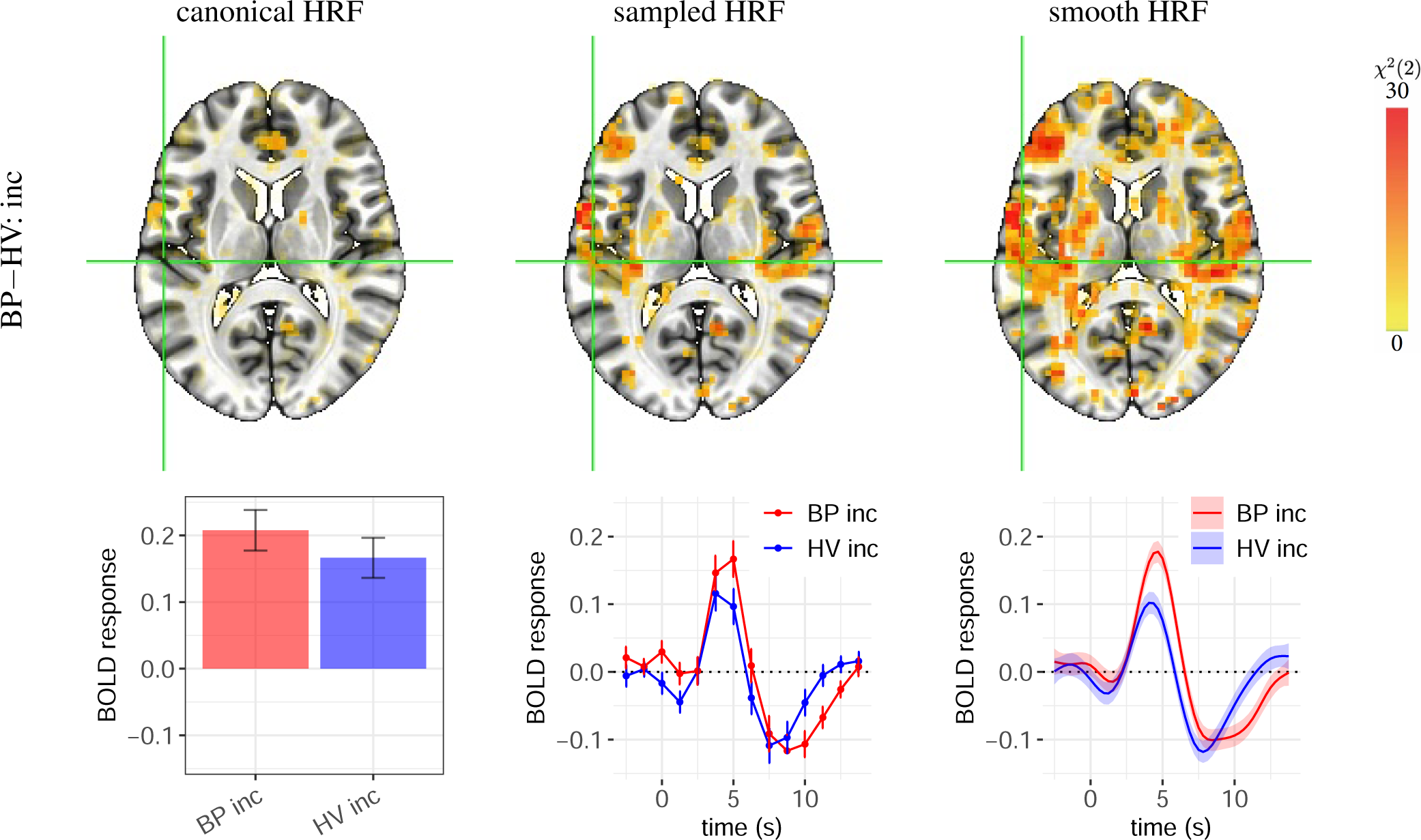
Case B: weaker statistical evidence by the canonical HRF compared to the HRF estimation approaches. The effect of focus is the group difference in HRF in the incongruent condition. The voxel at the cross-hair (*X,Y, Z*) = (56, 22, 11) is located in the left posterior insula. Vertical bars and shaded bands indicate standard error. As the estimated HRFs peaked close to what the canonical HRF assumes (i.e., 5 s), no underestimation of peak height occurred for either group, and the estimated group contrasts from the canonical HRF (first column) were roughly compatible with the HRF peak differences estimated via sampled and smooth HRFs. However, as the group difference (0.04 *±* 0.04) is much less than the case (0.15 *±* 0.03) in Fig. 5, the canonical HRF fails to show strong statistical evidence. In comparison, the characterization of the HRF shape through sampled and smooth HRFs provided strong statistical evidence for the presence of group difference. The other two HRFs (BP con, and HV con) and neighboring voxels share a similar HRF pattern (not shown here).

**Figure 7:**
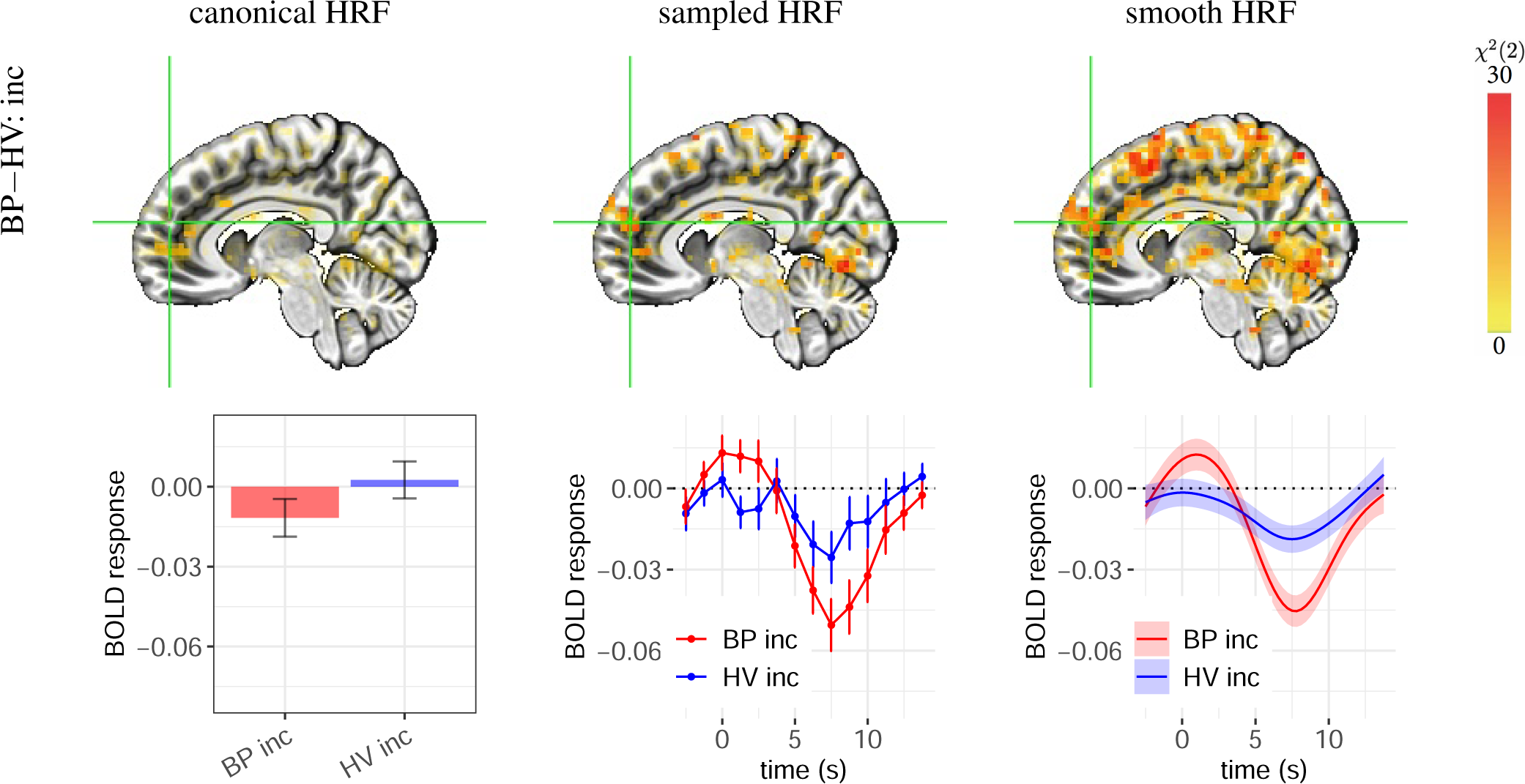
Case C: failure of the canonical HRF to capture BOLD responses resulting in an incorrect sign of effect estimation. The focal effect is the group HRF difference in the incongruent condition. The voxel at the cross-hair (*X,Y, Z*) = (*−*7*, −*52, 18) is located in the right medial frontal gyrus. Each vertical bar or shaded band indicates one standard error. The estimated HRFs had a small to negligible overshoot with a relatively large undershoot, which is dramatically different from the assumed HRF. Thus, the canonical approach failed to accurately capture the group difference in peak height. In comparison, morphological characterization through HRF estimation provided strong statistical evidence for the presence of a group difference. The other two HRFs (BP con, and HV con) and neighboring voxels shared a similar HRF pattern (not shown here).

**Figure 8:**
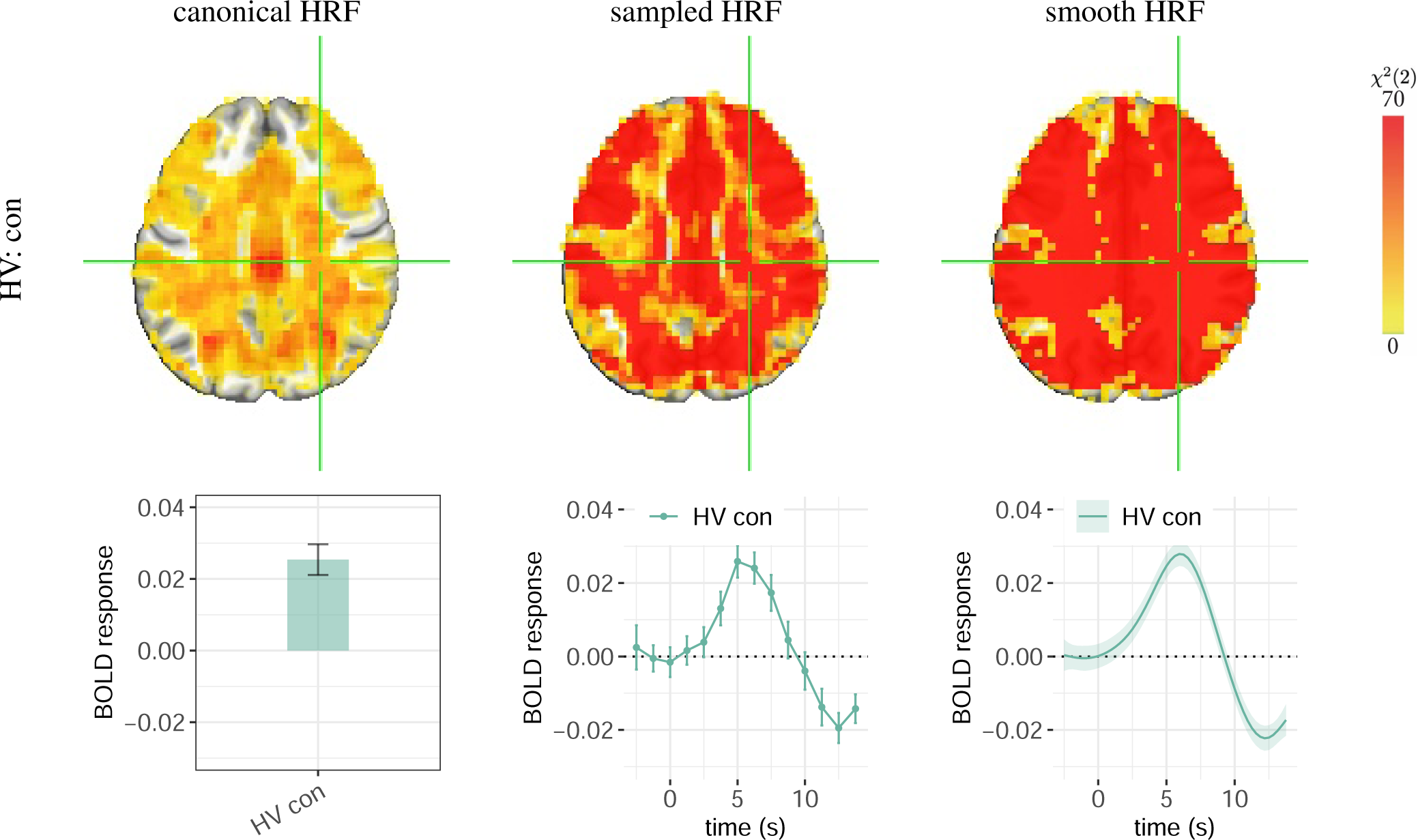
Statistical evidence for the presence of whole-brain BOLD response. The effect of focus is the congruent condition for the HV group. The voxel at the cross-hair (*X,Y, Z*) = (*−*28, 22, 32) is located in the right cerebral white matter on the axial slice. Each vertical bar or shaded band indicates one standard error. The BOLD response duration in white matter appeared to be longer than other regions in the brain. The other three HRFs (HV inc, BP con, and BP inc) and neighboring voxels shared a similar HRF pattern (not shown here). Strong evidence from both sampled and smooth HRFs indicates BOLD fluctuations cross the brain under the congruent condition. However, the canonical approach lacked sensitivity to detect effects in some regions (not colored) due to a close to zero or even negative magnitude estimate based on misaligned peak location and peak/nadir ratio. Fig. 9 shows several HRF shape properties for this same HV group’s congruent condition (see the fourth column for this *Z* = 32S slice); the associated peak magnitudes were typically nonzero but quite small in white matter.

**Figure 9:**
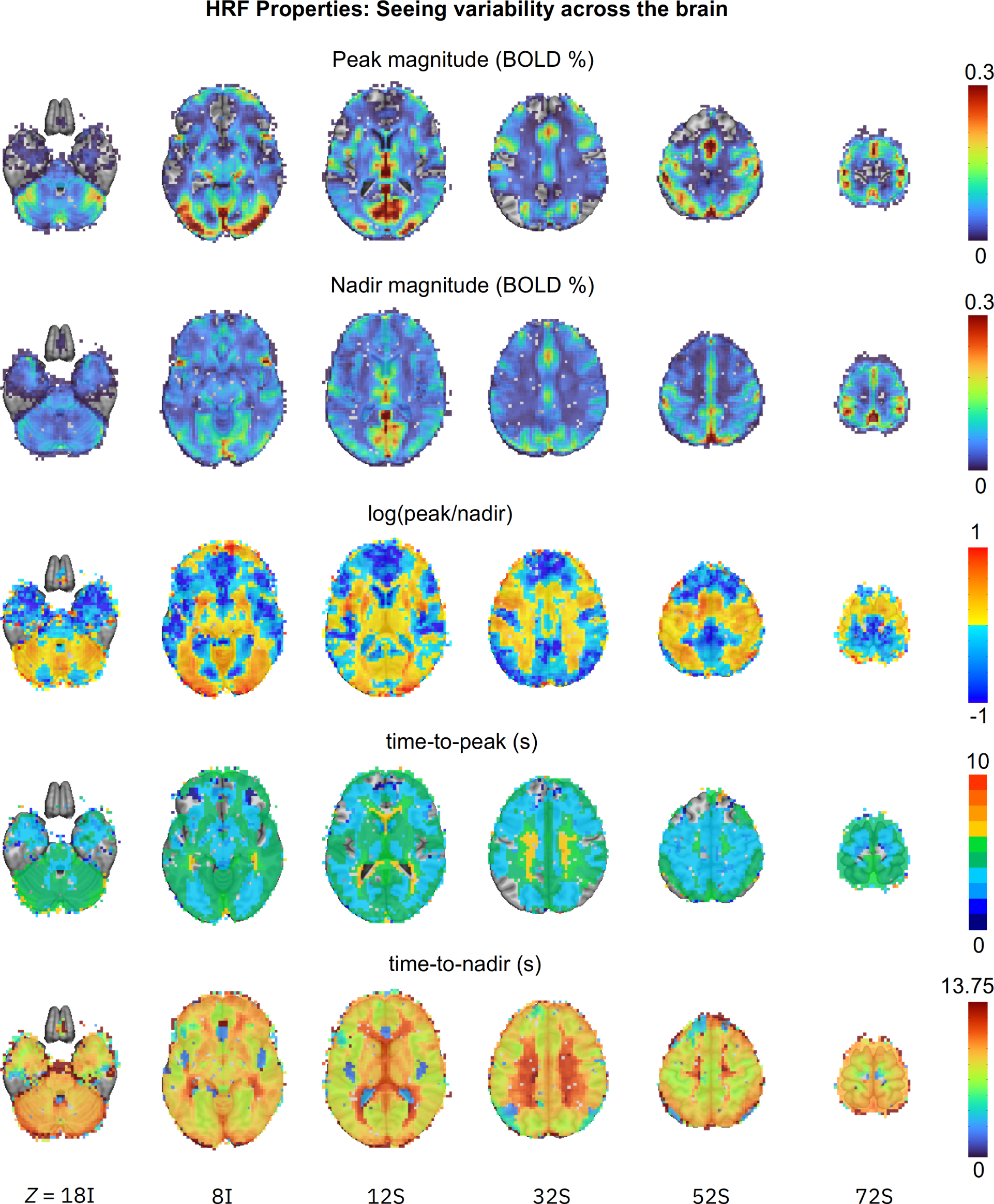
Examples of cross-brain variability of HRF shape features (six axial images, image left=brain left). Five key morphological properties are computed from the estimated HRFs at the population level and are shown for the HV group under the congruent condition with colors coding their respective magnitude: peak magnitude, nadir magnitude, log of the peak/nadir magnitude ratio, time-to-peak, and time-to-nadir. These quantities represent major features of interest in the HRF, and may not be exhaustive or exclusive; one might make analogy to diffusion tensor imaging (DTI), where interpretable scalar parameters of geometric features of interest are often examined (such as mean diffusivity, fractional anisotropy, etc.) in addition to the complete tensor set itself. “Empty” voxels in the first row show where the peak height was negligible, and in the second row, where the undershoot depth was close to 0. Several properties vary by an order of magnitude or more. Blue regions in the peak/nadir ratio panel show where the undershoot depth was greater than the overshoot peak. Non-green colors in the time-to-peak panel show where the actual HRF peak occurred more than 1 s away from the canonical HRF’s assumed 5 s.

We now explore some of the details which are observable with the smooth HRF approach, and particularly how this information would be misrepresented with the canonical HRF modeling. In each case, we explore how well different modeling approaches address the question, *Is there a difference in BOLD response between the BP and HV groups?* This will also reveal the breadth of response shapes which occur across the brain, further challenging the idea that a canonical HRF is representative of a global BOLD response shape.

**Case A.** Fig. 5 illustrates a location in the left postcentral gyrus that shows similarly strong statistical evidence in all three modeling approaches. However, when viewing the BOLD response estimates, one of the shortcomings of the canonical approach is evident. While each method shows a notable difference between BP and HV peak responses of approximately 0.15% signal change, it is clear that the peak responses from the canonical HRF are actually incorrect: the canonical approach assumes that the time to peak is *≈* 5 s, even though the overshoot actually peaked 1-2 s earlier. Therefore, BOLD responses in both groups were greatly underestimated. By happenstance, their difference *happens* to be equal to that at the actual peak. In comparison, both the sampled and smooth HRF methods provided more reasonable estimation.

This case shows that canonical HRF effect estimation, which relies strictly on shape feature assumptions and collapses all information to a single scalar, can result in substantial under-fitting and misestimation of the actual BOLD response. While the contrast may be correct in this case, this will not be the case in other instances. Indeed, as is already observable in the spatial maps in Fig. 5, the canonical HRF map has much weaker statistical evidence generally than those of the other two approaches. As most studies focus on contrasts between group/conditions, this would be particularly problematic in cases where the BOLD response either deviates from the canonical HRF only in one group/condition or deviates in different ways across groups/conditions, resulting in a systematically biased and inaccurate contrast estimates.

**Case B.** This example illustrates issues with the canonical approach, even if the assumed time-to-peak is largely accurate. In Fig. 6, the estimated HRFs peak around 4-5 s and have an overshoot shape similar to that of the canonical HRF. As a result, the peak estimates based on the canonical HRF (red and blue bars, Fig. 6) are roughly similar, compared to the peak estimates of the other two approaches (red and blue lines in Fig. 6). As a side note, the latter estimates also show that the undershoot appeared to be larger than that assumed by the canonical HRF, and the HV group had a noticeable initial dip. Despite the accurate capture of the peak height, the canonical approach did not provide strong statistical evidence, whereas the sampled and smooth HRF approaches did. This is likely due to several features, some of which are related to the collapse of so much fit information to a single scalar for the canonical HRF.

**Case C.** The third scenario showcases the failure of the canonical approach to detect group differences when the BOLD response shape differs substantially from the assumed shape. As shown in Fig. 7, each of the estimated group HRFs contained a small-to-negligible overshoot, but their undershoots vary greatly in magnitude. Because the morphology is so distinct from the canonical HRF, it is not surprising that this approach did not accurately capture the BOLD response magnitude, leading to a lack of strong statistical evidence for group differences in the incongruent condition. In fact, the canonical assumption of a peak at 5 s led to an incorrect sign (*−*0.012 *±* 0.007) for the BP group. The substantial deviation of the estimated HRFs from the standard shape was unlikely due to random noise, given the consistency of the HRF morphology between the two conditions (congruent and incongruent) within each group. Importantly, this non-standard HRF shape was not an isolated occurrence. Similar cases occurred in quite a few regions including the inferior, superior, central, and middle temporal gyri, precuneus, anterior cingular, and a number of regions that were predominantly white matter.

These cases exemplify three important points with regards to detection sensitivity and effect validation. First, the performance of the canonical HRF is highly sensitive to the peak location. When the underlying BOLD response is similar to the canonical HRF shape, the canonical approach can achieve strong statistical evidence. However, for voxels in the motor cortex shown in Fig. 5, the estimated HRF peaked around 3-4 s, slightly earlier than the canonical HRF and thus the peak estimates were substantially underestimated relative to the sampled and smooth HRF results. Second, the undershoot feature generally appeared to be larger than the one assumed in the canonical HRF. Such deviations of non-overshoot features can notably affect the overall HRF fit and therefore mischaracterize effect estimates. Third, there was also an indication of an initial dip for the HV group that was not captured by the canonical HRF. However, a higher temporal resolution would be required to investigate this feature in greater detail. Overall, while there is some convergence between the three approaches, these results also show that even relatively small deviations (e.g., an overshoot peak of 1-2 s earlier) from the rigid canonical shape can impact estimation accuracy—and below we see that this variability is indeed large. The much richer information conveyed through the estimated HRFs is important for both effect accuracy and statistical inference, as well as interpretability.

### 3.4 Evidence for BOLD fluctuations across the brain

In the previous subsection, we focused on a contrast of interest: BP-HV group difference for the incongruent condition. Here we focus on a single HRF to examine the question, *How much of the brain is involved during a simple task?* A previous study has investigated this question using BOLD responses averaged across many well-separated blocks, such as a simple visual stimulation or attention task with a deep (or dense) scanning paradigm with 100 runs in a block design for three participants (Gonzalez-Castillo et al., 2012). The results suggested that, as more runs were added, the BOLD response appeared to approach the whole brain level. Here, we investigate the same question using a lot less runs (but with many more subjects), as well as a fast event-related experiment. Compared to a block design, event-related designs usually have a much weaker effect magnitude, typically less than 1% signal change, due to the short stimulus duration. If this experimental paradigm would also support whole-brain fluctuations at the population level, it would provide an important piece of converging evidence for generalizability.

Fig. 8 shows that the sampled and smooth HRF estimation approaches provide strong evidence for population-level BOLD response throughout the majority of gray and white matter during the attention task. Overall, the BOLD response amplitude of the overshoot across the brain was largely below 0.1% signal change for either condition (see Fig. 9 for the HV group’s congruent condition). The HRF shape information offered evidence for the presence of BOLD response in nearly all regions, including white matter and cerebellum as well as those regions where the canonical approach failed to show strong statistical evidence (Fig. 8). These observations are consistent with previous findings in white matter (e.g., Gawryluk et al., 2014; Gore et al., 2019; Li et al., 2019). While the BOLD response in the cerebellum was of similar magnitude as in gray matter, its counterpart in white matter was at least a few times smaller (around 0.03% signal change for both the overshoot peak and undershoot nadir) with a longer duration (Figs. 8-9), consistent with a previous report (Li et al., 2019). Even though a similar HRF profile was also observed in the ventricles, it is much less clear how to understand this. Imperfect alignment and spatial smoothing could cause bleeding impact from neighboring brain regions, regardless of tissue type; these effects are particularly likely to be observed in small regions bordered by much larger ones. Nevertheless, we note that the evidence for the presence of typical HRF morphology occurred not only under one particular condition or group, but also across both conditions and both groups. The evidence of BOLD response is present not only in the white matter close to the gray matter, but also across the whole white matter (Fig. 8). In addition, some evidence exists in the literature for the involvement of neurovascular coupling driving the production and circulation of cerebrospinal fluid (Drew, 2022; Gonzalez-Castillo et al., 2022). We note that the evidence for the whole-brain fluctuations under a single condition is not inconsistent with other results regarding group/condition contrasts: little differential effect existed in their overall HRFs between groups or conditions; the magnitude of peak and nadir was quite small in most regions (Fig. 9).

### 3.5 HRF shape variations

Fig. 9 illustrates the spatial variability of HRF in detail by displaying key shape features throughout the brain for the HV group under the congruent condition (analogous inhomogeneity applies in all group/condition cases): peak magnitude, nadir magnitude, peak-nadir ratio, time-to-peak and time-to-nadir. In all cases, properties varied a lot. The overshoot peak was negligible or even absent in several parts of the frontal cortex (dark blue and empty overlay voxels in the top row, respectively). Recall that the canonical HRF assumes a fixed peak-nadir ratio of 9%; however, the ratio here had to be plotted with a logarithm because of the wide variability of values, and the peak magnitude is even less than the nadir magnitude in much of the brain (blue regions). The canonical HRF assumes a fixed time-to-peak of 5 s, but the actual times are mostly within a wider range the [4, 6] s interval, with a notable number of voxels in the lower range of [3, 4] s as well as a higher range of [6, 7] s, consistent with previous empirical individual-level results (e.g., Handwerker et al., 2004). Moreover, there does not appear to be a simple relationship among the shape properties or uniformity among tissue classes: these features reflect inherent shape differences.

In this dataset, there are many categories into which relative variability of HRF shape could be partitioned: between groups, between task conditions and across the brain. Overall, it was most pronounced spatially (Fig. 9), with some moderate degree of variation between groups and subtle variation between task conditions, in line with the images shown in Figs. 5-8.

## 4 Discussion

The contribution of FMRI studies to neuroscience heavily hinges on an accurate understanding of the link between the expected neural signal and the observed BOLD signal. For example, finer characterization of the BOLD response morphology will undoubtedly help capture cross-region relationships at the network level (e.g., Fouladirad et al., 2022; Gill et al., 2022). The underlying mechanism of neurovascular coupling remains to be fully understood - accurately characterizing the HRF morphology in modeling and effect estimation is a critical part of this endeavor. The stereotypical HRF is generally characterized with three temporal stages (Fig. 1): a transient initial dip, a hyperoxic peak, and a recovery undershoot. The explicit HRF estimation approach has been available at the individual level since the early days of neuroimaging data analysis, and all major software packages offer tools for implementation. However, these approaches are rarely applied in routine modeling, and even when they are adopted, their exploration usually either stops at the individual level or focuses on sampled HRFs. As we have shown here, there are many informative and quantitatively useful indices to more deeply explore the HRF shape in an analysis.

### 4.1 The importance of characterizing the HRF shape

The staple convolution methodology with an assumed HRF of neural events bears a strong assumption: the HRF shape remains the same across regions, individuals, tasks, conditions, and groups. It is this predetermined shape that allows the direct comparison of the BOLD response magnitude through a scalar and simplifies many modeling processes both at the individual and population level. This assumption makes neural decoding of FMRI data feasible through deconvolution (e.g., Uruñuela et al., 2022). Even when the assumption is moderately violated (e.g., a mischaracterized undershoot, absence of the initial dip), one would still be able to statistically detect the effect as long as its magnitude is relatively large and the focus is on contrasts rather than individual effects.

Here we show that there are many scenarios in which the canonical approach is less sensitive in detecting BOLD response. The consequences are under-fitting, effect distortion and compromised detection sensitivity, which likely contribute to reproducibility issues. Fig. 9 catalogues examples of observed HRF feature variation, including peak/nadir location and relative magnitude between peak and nadir. Importantly, we have shown several scenarios where the canonical approach underestimates (Fig. 5) or distorts (Fig. 7) the effect magnitude to varying degrees due to information reduction (Fig. 6) or shape deviation (Fig. 7). Most studies focus on a contrast between two conditions or groups, i.e. the difference between two scalars. Under the canonical approach this is achieved through two steps of information reduction: one from one-dimensional curves to zero-dimensional scalars, and the other from two such scalars to one difference value for the contrast. With no availability of information on the HRF shape, one is simply left with the sign of each scalar and cannot verify the accuracy of overshoot height, undershot depth or the combination of both.

One popular improvement is to adjust the canonical HRF, by either adopting one or two auxiliary bases or using a few shape metrics such as peak height, onset latency, time to peak and FWHM (Lindquist et al., 2009). However, while adjustments with auxiliary bases provide flexibility on location and width of the overshoot, these are not enough to capture possible profile variation, especially the undershoot variation and initial dip as shown in Figs. 5-8. Thus, their potential improvements remain limited (Chen et al., 2015), and the applicability of these metrics is difficult to model at the population level. For instance, simultaneously assessing these multiple features at the population level is challenging. In addition, the meaning of BOLD response magnitude may become ambiguous considering the following three different scenarios: a dominant overshoot, the coexistence of an overshoot and an undershoot, and an undershoot only. Additionally, when the auxiliary bases are used to adjust the peak location and width of the overshoot, despite improved model fit, the effects associated with such auxiliary bases are usually then ignored, resulting in effect accuracy issues (Figs. 5,7). Even with available tools, it remains a challenge to effectively incorporate the overshoot adjustments into follow-up analysis. Furthermore, a canonical HRF is intended to capture an overshoot that peaks around 5 s, and an undershoot which is fixed and small (9% of the overshoot peak). As a result, the likelihood of accurately capturing an undershoot becomes slim – with the added complication that sometimes the undershoot is even more dominant (Fig. 7).

### 4.2 Advantages of estimating HRF shapes

The sampled HRF approach is rarely used in practice. While it has clear advantages, such as increased sensitivity, there are also some disadvantages with sampled HRFs that need to be discussed. First, more regressors would be required to capture the HRF shape. In other words, HRF estimation approaches require careful experimental designs to avoid collinearity issues. Furthermore, without constraints imposed on the HRF shape, the basis functions may catch signal unrelated to BOLD response or suffer from sampling variations, as evident by the jagged appearance of estimated HRFs even at the population level through sampled HRFs (Figs. 5-8). Lastly, it is a computationally cumbersome task to estimate HRFs at the population level, which may be the main reason for why this approach is rarely attempted.

Despite its improvements relative to previous methods, the sampled HRF approach (Chen et al., 2015) needs further refinement. Due to sampling errors, HRF estimates even at the population level still contain zigzags, a common symptom of over-fitting (second column in Figs. 5-8). In other words, when one fully trusts individual-level HRF samples, it is unavoidable to capture some noise. In contrast, even though HRF estimates at the individual level are purely data-driven and sampled at a temporally coarse resolution, the estimation through smooth splines at the population level balances between under- and over-fitting. Hence, we make the basic and reasonable assumption that the HRF should be smooth, i.e., not fully fitting to the raw data. Yet, we still learn the HRF features and profiles from the data.

To summarize our results, the performance of the three modeling approaches at the population level follows the order to which they reduce information: the sampled HRF approach is more accurate and sensitive than its canonical counterpart, but it does not perform better than the smooth version. Below are a few specific advantages of HRF estimation.

a. Improved detection sensitivity. Modeling in neuroimaging comprises two intertwined components: causal inference (e.g., group or condition differences) and predictive accuracy. While our data indicate that the HRF is very close to the canonical curve in motor and visual regions, the peak in those two regions occurred 1-2 s earlier than is stipulated by the canonical HRF. This small deviation leads to substantial distortion in effect estimates. While a more complex set of models, HRF estimation achieved both enhanced accuracy and sensitivity. The improved sensitivity is particularly more valuable for higher-order effects such as interactions than main effects due to the former’s requirement for much larger sample sizes (Leon and Heo, 2009; Gelman et al., 2020).
b. Improved reproducibility. As effect magnitudes are rarely reported in the literature (especially for voxel-wise analyses), neuroimaging analyses often have many “experimenter degrees of freedom” which contribute to reproducibility issues (e.g., Chen et al., 2022b). When HRFs are directly estimated, the detailed information (i.e., HRF profiles) can be utilized to assess the validity of an effect as an extra piece of confirmatory information, independent of the statistical evidence. This approach promotes a shift from a decision-making process “blind” to the data but only based on statistical evidence, to focusing on effect estimation and morphological subtleties. For example, the current practice of handling multiplicity solely through statistical evidence in the spatial neighborhood could be improved: instead of making a dichotomous decision, we propose a holistic approach to providing more compelling information than statistical evidence alone. Such information can be the combination with other sources of evidence including the presence of BOLD response in the form of HRF morphology, anatomical relatedness (e.g., HRF morphological symmetry between contralateral regions) as well as prior information (e.g., literature).
c. Exploring differences in HRF shape. Our investigations in the context of the sustained attention task indicate that HRF variability across regions and groups is much larger than condition differences. Even though not all HRF shape subtleties are directly associated with BOLD response magnitude, accurate characterization opens an avenue for future investigations regarding the underlying neurovascular structure and associated neurological and hemodynamic mechanisms. For example, some (1-2 TRs worth) of anticipation effects occurred in some regions (Fig. 5-7). It is generally recognized that BOLD effects include changes in cerebral blood flow, volume, metabolic rate of oxygen, and hematocrit fraction, and is further complicated by various vascular origins. Moreover, the undershoot, the mechanisms of which are poorly understood, may be due to a few factors such as delayed vascular compliance, sustained increases in cerebral metabolic rate of oxygen, post-activation reduction in cerebral blood flow (van Zijl et al., 2012), or inhibitory processes (Mullinger et al., 2017). Previously, little attention has been paid to undershoots in modeling. Now, their presence could be directly estimated and visually verified through HRF estimation. Similarly, the initial dip in BOLD signal is typically not considered important in modeling even though optical intrinsic signal imaging studies routinely observe its presence (Das and Gilbert, 1997; Chen et al., 2005). In addition, an initial dip can be predicted by quantitative models that characterize the relationships among blood flow, metabolic rate of oxygen, and BOLD signal (Buxton et al., 2004; Maith et al., 2022). Here, we empirically demonstrated that through HRF estimation, an initial dip was seen in some brain regions (Fig. 6) even though a finer temporal resolution would be required to improve certainty.

Questions exist on the degree to which estimated HRFs can be used to make inferences at the network level? As the BOLD response shape varies across individuals, regions, groups and task conditions, our current modeling focus is limited to region level inferences through univariate analysis. Extensive work reveals networks that are engaged during various task conditions through multivariate modeling using constrained principal component analysis with estimated HRFs (e.g., Woodward et al., 2016; Sanford and Woodward, 2021; Roes et al., 2022). In these studies, dominant networks emerge based on similarity in HRF shape. Univariate and multivariate approaches adopt different assumptions and possess different goals. The hemodynamic response shape is only a proxy – not an exact representation – of neural activity. Accordingly, cross-regional inferences based on response morphology may lose some sensitivity and accuracy given neurovascular heterogeneity across regions. Therefore, univariate modeling is likely more sensitive in detecting subtle differences across groups and task conditions at the region level, differences which multivariate approaches may have difficulty identifying. However, inferences based on univariate analyses are largely confined to the local level without integration at the network level. Future work could explore how these two approaches complement each other.

Now we can directly address the questions raised in the Introduction.

1. *How much does the HRF shape vary across space, tasks, conditions, and groups?* While there is variability in each category, the observed shape variability was by far the largest across the brain. This is consistent with the reports of cross-region heterogeneity in draining veins: regions with larger draining veins showed more delayed BOLD response (Handwerker et al., 2004; Taylor et al., 2018a), and the variability was also large in the rest of the brain. Some subtle variations occurred between clinical groups, with a similar profile across conditions.
2. *Does the smooth HRF approach improve effect detection sensitivity and efficiency?* Yes, the modeling approach with smooth splines captured the HRF shape subtleties and reduced information loss compared to the traditional approaches (canonical and sampled HRFs).
3. *Does the HRF shape offer validation support for statistical evidence?* Yes, we believe that visual verification of the HRF shape can play a confirmatory role, independent from the statistical evidence, and verify the likelihood of effect underestimation and incorrect signs by the canonical approach.
4. *Is the whole brain involved during a simple task?* Through capturing the HRF shape with an event-related experiment, our dataset provided evidence for whole-brain response at the population level, and these results are similar to what was revealed previously at the individual level via an experiment of deep scanning (Gonzalez-Castillo et al., 2012).
5. *Does the HRF estimation approach provide evidence for BOLD response in white matter?* Yes, our results revealed hemodynamic activities with the HRF signature shape yet with small peak magnitude.

Future explorations are required to further validate the modeling approach with smooth splines. The improvements in model quality at the population level, as assessed in questions 1) and 2) above, have been established, to some extent at the individual level in previous work. The increased detection sensitivity by smooth HRFs relative to sampled HRFs, as demonstrated in Figs. 4-8, is notable but not as dramatic compared to the canonical HRF except for the interaction (third row, Fig. 4). Subsequent studies are needed to evaluate the strength of the smooth HRF in its predictive accuracy when applied to out-of-sample data; here, we have only validated the modeling approach in a single case study.

This work provides new insights and extensions to the common practice in the field of FMRI. The modeling approach with a canonical HRF has been and will likely continue to play a crucial role in neuroimaging. However, its inevitable imprecision may be a contributor to reproducibility issues faced by the field. For example, the adoption of a canonical HRF likely has ramifications for the challenge of estimating intraclass correlation or test-retest reliability (Chen et al., 2021b). Since psychophysiological interaction (Friston et al., 1997) analyses also rely on deconvolution using a canonical HRF, many of the points we raise here may also be applicable. Moreover, the large cross-region heterogeneity of BOLD response morphology presents a challenge: the sensitivity for cross-region relationships (e.g., correlation) that are heavily based on the similarity of response shapes could be compromised. The omnipresence of BOLD response in the brain also has implications. Our observations of HRF morphology also confirmed previous results suggesting a smaller overshoot peak, delayed onsets and prolonged initial dips in white matter compared to gray matter (Li et al., 2019). Lastly, Moroeve, it raises the question regarding the common practice of using the average signal in white matter as a regressor of no interest in resting-state data preprocessing (Jo et al., 2013; Power et al., 2014; Ciric et al., 2017), and echoes the concern about the blind spot status of white matter (Grajauskas et al., 2019).

### 4.3 Limitations of HRF estimation through smooth splines

HRF estimation can be considered an extension to linear regression: instead of fitting the data with straight lines, we model the data with nonlinear curves. Just as the linearity assumption in a traditional regression model can be violated to a varying degree or face other issues such as distribution misrepresentation, so do limitations and challenges exist for HRF estimation through smooth splines.

1. The appropriateness of the smoothness assumption versus temporal resolution. The temporal smoothness assumption for the HRF (up to second-order derivatives) is likely reasonable most of the time. However, it is possible that some underlying shape subtleties may require a higher-than-available temporal resolution when acquiring the data, in order to be able to capture features accurately. Violating this assumption could lead to over-smoothing or under-fitting. For example, without an adequately short TR, it may become difficult to accurately capture the initial dip and overshoot peak height/location.
2. Complexity in experimental design. To avoid potential collinearity, one mush carefully design an experiment so that the model can accurately estimate the HRF shape information at the individual level. Such designs may need to carefully consider the use of randomization to reduce collinearity through variations in inter-trial intervals and jittering of event timings across conditions. Furthermore, a short TR (e.g., 2 s or less) may be required to reduce the risk of under-fitting.
3. Complications in modeling and result reporting. Despite its flexibility of capturing HRF profiles, the approach still requires a prior determination of the BOLD response duration in individual-level modeling. For example, even though a duration of 13 TRs (i.e., 16.25 s) was enough to cover the responses at most regions with our experimental dataset, data suggest that a longer duration is required in white matter (Fig. 8). Future work is needed to understand how individual-level measurement errors could be incorporated into the population-level HRF estimation. Moreover, it remains challenging to estimate trial-level HRFs, which is important for statistical learning processes utilized in multi-voxel pattern analysis, support vector machine and linear discriminant analysis. In addition, detection sensitivity comes at a cost: unlike a single scalar for a canonical HRF that can directly reveal the sign of a comparison (positive or negative), each estimated HRF is expressed as a full curve.

Therefore, one would have to resort to visual inspection to assess the relationship across conditions or groups, presenting a challenge for result reporting as well as for meta-analyses across studies.

The modeling approach with smooth splines is summarized in Table 3. The current work leaves some issues unresolved. The association of the overshoot with the excitatory process of neural activity, is the theoretical underpinning of a canonical HRF approach. However, it remains unclear whether the overshoot peak should be the sole focus irrespective of the undershoot magnitude. Our data indicates that the relative magnitude of the overshoot ranges from larger to smaller than the undershoot nadir across the brain; the associated mechanism for neurovascluar coupling is opaque. In addition, HRF variability across individuals and across trials remains to be explored in future work.

**Table 3:** Comparisons of properties among different HRF modeling approaches.

| Individual level |  |  |  |
| --- | --- | --- | --- |
|  | Canonical HRF | Sampled HRF | Smooth HRF |
| shape assumption: | (a priori) fixed | adaptive, data-driven |  |
| basis: | Gamma variates | deconvolution with piecewise linear splines (e.g., TENT, FIR) |  |
| dimension: | scalar (0D) | vector (1D) of equally spaced samples |  |
| trial-level effects: | possible | difficult |  |
| Population level |  |  |  |
|  | Canonical HRF | Sampled HRF | Smooth HRF |
| dimension: | scalar (0D) | vector (1D) of equally spaced samples | smooth curve (1D) |
| interpretability: | high: signed peak magnitude | moderate: detailed shape features, but relies on visual examination |  |
| robustness: | low: inflexible and vulnerable to under-fitting | moderate: adaptive but vulnerable to over-fitting | high: adaptive and regularized |

### 4.4 Suggestions for experimental designs and modeling

Based on this work, we would make the following recommendations for researchers designing task-based FMRI studies with the intention of implementing HRF estimation in data analysis:

1. The importance of careful experimental design cannot be overemphasized. Estimating HRFs at the individual level is a crucial first step in the procedure, and it necessitates many more regressors than the canonical approach (e.g., one per TR within the expected stimulus duration). Thus, both inter-stimulus jittering and trial sequence randomization across conditions are critical to avoid potential collinearity across events. In addition, to improve statistical efficiency, having a reasonable trial sample size is almost as important as the participant sample size (Chen et al., 2022a).
2. Having a higher temporal resolution helps to detect subtle differences in HRF shapes. Because of the potentially complicated nonlinearity involved in the HRF shape, one needs enough basis functions to capture the fine structure at the individual level (and the maximum number of basis functions is limited to the number of TRs within the HRF interval). We suggest using seven bases as a minimum. With a TR of 2 s, this translates to 12 s of HRF duration. A small TR (e.g., less than 2 s) would likely improve the accuracy of shape characterization, e.g., for localizing the peak and detecting an initial dip, among other features. Alternatively, one may improve the temporal resolution through leveraging the participant sample size. For example, even with a TR of 2 s, one could collect HRF samples at the TR grids for half of the subjects, and obtain shifted samples by 1/2 TR (i.e., 1 s) for the other half. In doing so, HRFs could be estimated at a finer temporal resolution of 1 s at the population level.

Modeling advances will continue to play an important role in improving neuroimaging data analysis. FMRI signal is “contaminated” with many unknown components that currently remain unaccounted for. In fact, the amount of BOLD variability accounted for in models remains quite low: 10% or less for event-related experiments in most brain regions among typical condition-level analyses. When statistical evidence fails to reach a designated level, there is often a strong tendency to ascribe the failure to a small effect size or suboptimal sample sizes. However, attempting to remedy this by simply increasing the number of samples, although potentially increasing statistical evidence could be inefficient and costly.

## 5 Conclusions

In this FMRI study, we observed that a substantial amount of HRF shape variability occurred across the brain and that the canonical HRF assumptions are often poorly suited to modeling it. As a consequence, some subtle differences between groups and conditions, which are typically the focal point of such studies, can be lost or misrepresented. To help address this issue, we introduced a modeling approach of estimating HRFs at the population level, modeling nonlinearity through smooth splines. With the simple assumption that the BOLD response is likely to be smooth rather than rough, the hemodynamic profile can be effectively estimated without knowing a priori the specifics of its shape characteristics. The process of uncovering the HRF effects through smooth modeling becomes relatively robust when the time resolution is fine enough. Furthermore, the validity of estimated smooth HRFs can be visually verified and compared across groups and conditions. We hope that the modeling framework, with the associated program 3dMSS, will contribute to increasing effect detection sensitivity and to improving reproducibility in neuroimaging data analysis.

## Acknowledgments

The research and writing of the article were supported (GC, PT and RCR) by the NIMH Intramural Research Program (ZICMH002888) of NIH / HHS, USA. Data collection was supported (EL) by the NIMH Intramural Research Program (ZIAMH002786) and was carried out under the National Institutes of Health Clinical Study Protocol 00-M-0198 (ClinicalTrials.gov ID: NCT00006177). This work used the computational resources of the NIH HPC Biowulf cluster (http://hpc.nih.gov). Additional support for SH was provided by a BBRF Young Investigator Grant.

## Footnotes

1 The undershoot or negative BOLD response is often referred to as “post-stimulus”, a term likely originated from experiments with block designs, in which the undershoot occurs after the end of each stimulus presentation. In event-related designs, its usage becomes obscure, because both initial dip and overshoot (or positive BOLD signal), if present, also occur after stimulus onset.

2 Throughout this work, the following notation for mathematical expressions is used: regular, lowercase italic letters (e.g., *b*) stand for scalar parameters or variables; boldfaced, lowercase italic letters (e.g., *b*) for column vectors; and boldfaced, uppercase italic letters (e.g., *B*) for matrices.

